# *Toxoplasma gondii* actin filaments are tuned for rapid disassembly and turnover

**DOI:** 10.1101/2023.08.29.555340

**Authors:** Kelli L. Hvorecny, Thomas E. Sladewski, Enrique M. De La Cruz, Justin M. Kollman, Aoife T. Heaslip

## Abstract

The cytoskeletal protein actin plays a critical role in the pathogenicity of *Toxoplasma gondii*, mediating invasion and egress, cargo transport, and organelle inheritance. Advances in live cell imaging have revealed extensive filamentous actin networks in the Apicomplexan parasite, but there is conflicting data regarding the biochemical and biophysical properties of *Toxoplasma* actin. Here, we imaged the *in vitro* assembly of individual *Toxoplasma* actin filaments in real time, showing that native, unstabilized filaments grow tens of microns in length. Unlike skeletal muscle actin, *Toxoplasma* filaments intrinsically undergo rapid treadmilling due to a high critical concentration, fast monomer dissociation, and rapid nucleotide exchange. Cryo-EM structures of stabilized and unstabilized filaments show an architecture like skeletal actin, with differences in assembly contacts in the D-loop that explain the dynamic nature of the filament, likely a conserved feature of Apicomplexan actin. This work demonstrates that evolutionary changes at assembly interfaces can tune dynamic properties of actin filaments without disrupting their conserved structure.

## Introduction

The human pathogen *Toxoplasma gondii* (*T. gondii*) causes life-threatening disease in immunocompromised individuals and when infection occurs *in utero* (1–3). *T. gondii* is a member of the Apicomplexan phylum that encompasses over 5000 species of parasites including a large number of parasites of medical and veterinary importance, including *Cryptosporidium spp*. and *Plasmodium spp*., the causative agents of cryptosporidiosis and malaria respectively.

*T. gondii* is an obligate intracellular parasite. Survival and disease pathogenesis are dependent on host cell invasion, intracellular replication and egress, which results in destruction of the infected cells. *T. gondii* expresses a single divergent actin gene (*TgAct1*) that is essential for successful completion of this lytic cycle. This isoform shares only 83% similarity to skeletal alpha actin and mammalian β and γ isoforms (4) and was originally shown to be essential for the gliding motility of the parasite, host cell invasion and egress (5–8). More recent studies have greatly expanded our understanding of other essential functions of actin in intracellular parasites. These include: inheritance of a non-photosynthetic plastid organelle named the apicoplast (9, 10), the directed movement of secretory vesicles called dense granules (11), recycling of secretory vesicles called micronemes in dividing parasites (12), morphology and positioning of Golgi and post-Golgi compartments, the movement of ER tubules (13) and conoid extension (14).

Despite its ubiquitous functions in Apicomplexan parasites, the organization of F-actin in *T. gondii* remained elusive for many years because conventional actin probes such as phalloidin, GFP-tagged actin, and LifeAct failed to detect filamentous structures (15). The absence of filamentous actin was further supported by biochemical studies which showed that recombinantly purified HIS-TgAct1 formed only short unstable filaments *in vitro* (reviewed in (15)). This also seemed to be the case for the closely related *Plasmodium falciparum* actin 1, which shares 93% sequence identity with TgAct1.

The idea that Apicomplexan actin does not polymerize into long filaments *in vivo* has recently been challenged with the development of an actin chromobody as a tool for imaging F-actin in *T. gondii* and *P. falciparum* (16–18). Using this approach, studies have revealed vast tubular actin networks that connect individual parasites within host cells and remarkably dynamic actin networks within the parasite cytosol (17, 18). The success of using the actin chromobody is likely because it shows minimal effects on actin dynamics compared to other probes (19, 20). Recently, recombinantly purified actin chromobody has been used as a tool to study the dynamics of recombinantly purified *P. falciparum* actin *in vitro* and showed that it can polymerize into long dynamic filaments (21). These results are more consistent with the observation of filamentous networks in the cell and also indicate that assembly factors are not required for its polymerization.

While these studies have provided novel insights into TgAct1 organization, many biochemical and structural questions remain. There is conflicting data on whether TgAct1 has a high or low critical concentration, as both have been reported for Apicomplexan actin (21–23). It has also been reported that TgAct1 filaments assemble isodesmically, and therefore lack a critical concentration (22). Intriguingly, long filaments of TgAct1 or *P. falciparum* Act1 have rarely been observed directly using electron microscopy without stabilizing agents, which has led to the idea that long filaments of TgAct1 can only form in the presence of stabilizers (24). A lack of high-resolution structures of unstabilized TgAct1 or *P. falciparum* Act1 filaments has also hindered exploration of the unusual properties of Apicomplexan actin.

In this work, we used epifluorescence microscopy to image the growth of TgAct1 filaments *in vitro*. We find that TgAct1 can form filaments tens of microns in length that rapidly treadmill. This feature is due to an unusually rapid subunit dissociation constant at the pointed-end and a nucleotide exchange rate constant from the monomer that is 50-fold faster than that of skeletal muscle actin. Using electron cryomicroscopy, we determined the structures of both jasplakinolide-stabilized and native, unstabilized TgAct1 filaments. In these two conditions, we found that the DNase I-binding loop (D-loop) adopts different conformations, with the unstabilized conformation likely contributing to rapid filament subunit dissociation. Taken together, this study not only resolves outstanding questions in the field regarding the ability of TgAct1 to form long filaments and its assembly properties, but also demonstrates how new dynamic properties can evolve within the constraints of a filament architecture.

## Results

### TgAct1 polymerizes into long filaments in vitro with a high critical concentration

Because modifications to actin, such as tags or direct labeling, can influence the biochemical and kinetic properties of actin polymerization, we purified unmodified TgAct1. Untagged TgAct1 (ToxoDB: TgME49_209030) was prepared by expressing the gene in SF9 cells fused to a C-terminal β-thymosin–6xHIS tag, which maintains the actin in a monomeric state during expression and allows for removal by chymotrypsin, resulting in an untagged and unmodified protein product (**Fig. 1a and b, Ext. Data Fig. 1a**). This strategy was previously used for the expression of *P. falciparum* and *Dictyostelium* actin, which shares 95% and 85% identity to TgAct1 respectively (21, 25, 26).

**Figure 1:**
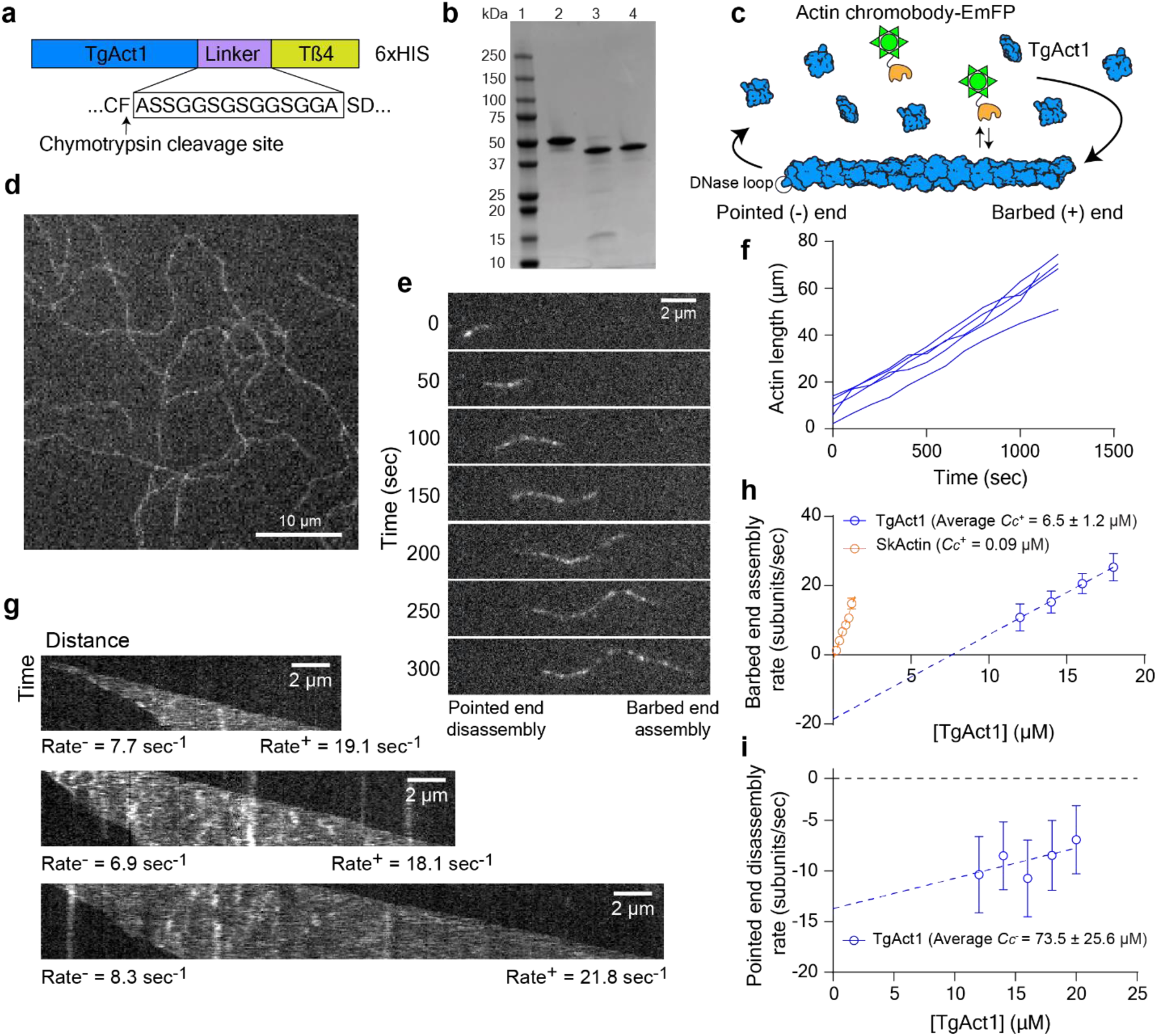
Imaging and analysis of TgAct1 polymerization *in vitro*. **a**, Domain organization of TgAct1 fused to a β-thymosin–HIS tag. After expression and purification from Sf9 cells, the tag was cleaved by chymotrypsin after the terminal phenylalanine producing native, untagged TgAct1. **b,** Coomassie stained SDS-PAGE gel of (lane 1) protein molecular weight marker; (lane 2) TgAct1–β-thymosin–HIS following HIS purification; (lane 3) TgAct1 after chymotryptic digest; (lane 4) purified TgAct1 after ion-exchange and size exclusion chromatography. **c,** Schematic of the *in vitro* polymerization assay. TgAct1 monomers (blue) were induced for polymerization and added to a blocked flow chamber absorbed with NEM-myosin II to maintain a semi-stable attachment for imaging. Epifluorescence microscopy was used to image the TgAct1 filament dynamics by inclusion of 25-50 nM actin chromobody fused to EmeraldFP (EmFP) (green), which interacts transiently with the growing filaments. **d,** Static epifluorescence microscopy image of TgAct1 filaments in the *in vitro* polymerization assay showing filament lengths > 60 µm. **e,** Montage of a single TgAct1 filament that is shrinking from the pointed (-) end and growing from the barbed (+) end in the presence of 16 µM TgAct1 monomers. Blue dashed line indicates the position of the (-) end of the filament at time zero. **f,** Kymographs for three individual filaments in the presence of 16µM TgAct1 monomers showing disassembly from the pointed (-) end (left side) and assembly from the barbed (+) end (right side). Rates for individual filaments are shown in subunits per second. **g,** Measuring the filament length of 5 representative filaments over time shows constant growth rate over 1200 sec of imaging. **h,** Plot of the rate of barbed (+) end growth in subunits/sec, per actin concentration for TgAct1 (blue) compared to skeletal actin (orange). Data shown is an aggregate of three independent preps. The average critical concentration (*C_c_*), determined by the x-intercept of the fitted line, is higher for TgAct1 (6.5 µM) compared to skeletal actin (0.09 µM). **i,** Plot of barbed (-) end disassembly rate per TgAct1 concentration. The average *C_c_* for the barbed end, determined by the x-intercept of the fitted line, is 73.5 µM. Imaging conditions: 25 mM imidazole, pH 7.4, 50 mM KCl, 2.5 mM MgCl_2_, 1 mM EGTA, 2.5 mM MgATP,10 mM DTT, 0.25% methylcellulose, 2.5 mg/mL BSA, 0.5% Pluronic F-127, oxygen scavenging system (0.13 mg/mL glucose oxidase, 50 μg/mL catalase, and 3 mg/mL glucose), 37°C.

Polymerization of individual actin filaments was imaged in real-time using an epifluorescence microscopy-based *in vitro* actin polymerization assay (**Fig. 1c**). The fluorophore, a recombinantly purified actin chromobody tagged with EmeraldFP (**Ext. Data Fig. 1b**, hereafter referred to as chromobodies) was previously used to observe filament dynamics and organization in *T. gondii* cells (16), and as a novel method for imaging *in vitro* polymerization of *P. falciparum* actin (21). The advantage of this approach is that actin polymerization can be visualized without fluorescent labeling of actin monomers, which has been shown to dramatically affect skeletal actin assembly and disassembly rate constants (27). Consistent with this, we find that TgAct1 conjugated directly with rhodamine to Cys374 shows poor incorporation into filaments (**Ext. Data Fig. 1c**), and thus unsuitable for visualizing TgAct1 polymerization *in vitro*.

Using the chromobody approach, we observe robust TgAct1 polymerization at concentrations above 12 µM actin, with filaments growing to greater than 60 µm in length (**Fig. 1d-f and Video S1**). To demonstrate that TgAct1 can form long filaments in the absence of the chromobody, we polymerized filaments in a flow chamber without chromobodies for 10 minutes. Chromobodies were then added and imaged before filaments disassembled in the absence of actin monomers. When we do this, we observe filaments longer than 10 µm, indicating that TgAct1 is able to polymerize into long filaments in the absence of chromobodies or other stabilizing agents such as jasplakinolide (**Ext. Data Fig. 2)**.

**Figure 2:**
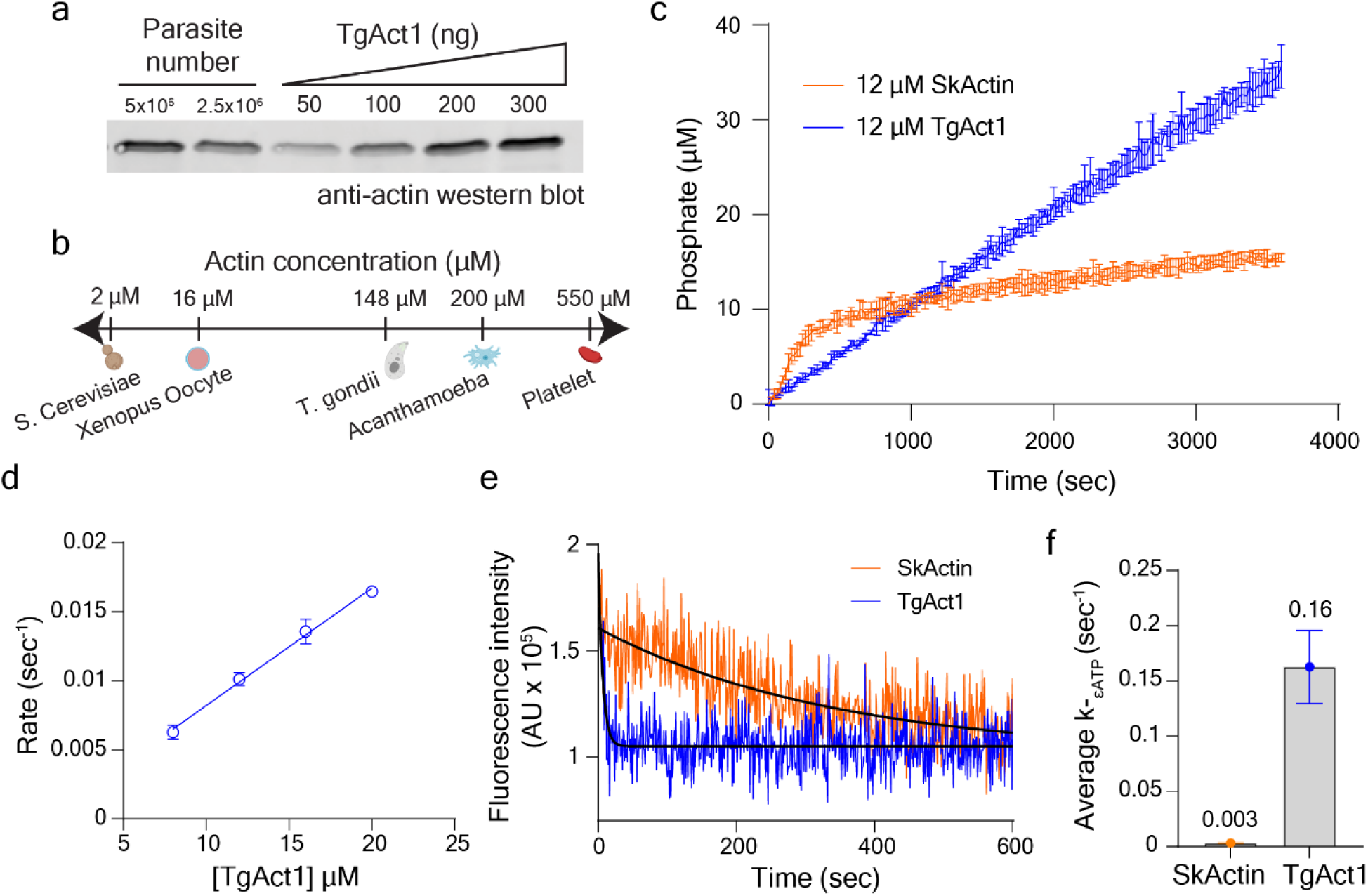
Cellular concentration and kinetic analysis of TgAct1. **a**, anti-actin western blot of *T. gondii* parasite lysate and known amounts of purified TgAct1. The average cellular concentration of TgAct1 (148 ± 7.6 µM) was determined by comparing the relative band intensity of a known number of parasites to those of known amounts of purified TgAct1. Error ± SEM, n=4. **b,** Comparison of TgAct1 cellular concentrations to other known model systems (79). Cartoons created with BioRender.com. **c,** Time course of phosphate release from 12 µM TgAct1 (blue) and skeletal actin (orange) after inducing polymerization. **d,** Plot of ATP hydrolysis rates over a range of TgAct1 concentrations. Conditions for ATPase and ɛATP experiments: 25 mM Imidazole, pH 7.4, 50 mM KCl, 1 mM EGTA, 2 mM MgCl_2_, 0.2 mM MgATP,1 mM DTT, 37°C. **e,** Plot used to determine the dissociation rate constant (*k*_-ATP_) of ATP for TgAct1 (blue) and skeletal actin (orange). Fluorescence decay of ɛATP-bound actin after addition of a large molar excess of MgATP was measured over time. *k*_-ATP_ was determined by exponential fit to the data. Conditions: 20 µM actin, 25 mM Imidazole, pH 7.4, 50 mM KCl, 1 mM EGTA, 2 mM MgCl_2_,1 mM DTT, 37°C. **f,** Bar graph of average *k*_-ATP_ for TgAct1 (blue, 0.16 ± 0.03) and skeletal actin (orange, 0.003 ± 0.0002). Error ± SD, n=3.

Strikingly, over the concentration range measured (12-24 µM), individual filaments treadmill, depolymerizing at the pointed-end while elongating at the barbed-end (**Fig. 1e and Video S2**). Treadmilling is readily visualized in kymographs (distance vs time), which shows pointed-end disassembly (left) and barbed-end elongation (right) of representative filaments at an actin concentration of 16 µM (**Fig. 1g**). This property is also a characteristic of *P. falciparum* actin (21) and is thus a likely conserved feature of Apicomplexan actins, which are all divergent from skeletal muscle actin (4, 28). Long term imaging of actin polymerization shows that filaments continue to polymerize at a constant rate for 20 minutes of imaging (**Fig. 1f**).

The actin concentration-dependence of assembly and disassembly rates provides the critical concentration (*C*_c_) and the rate constants for subunit incorporation and dissociation at the barbed and pointed ends of filaments. A linear fit to plots of the assembly or disassembly versus actin concentration can be used to determine the critical concentration (x-intercept), subunit dissociation rate constant (y-intercept) and subunit association rate constant (slope) at each end (29). As a control, we imaged the polymerization of different concentrations of skeletal muscle actin using the chromobody and found that the critical concentration (0.09 µM), disassembly rate constant (1.2 sec^-1^) and assembly rate constant (12.7 μM^-1^·s^-1^) are consistent with previously reported values (29), which is further evidence that the chromobodies do not influence the polymerization, critical concentration, or subunit incorporation rate constants (**Fig 1h, orange, Table 1**).

**Table 1:** Summary of polymerization rate constants for TgAct1 and skeletal actin. Rate constants for the barbed (+) and pointed (-) end of TgAct1 and skeletal actin were determined from plots of assembly (barbed-end) or disassembly (pointed-end) rates versus actin concentration. The critical concentration (*C_c_*) is defined by the x-intercept. The assembly (*k*_+_) and disassembly (*k*_-_) rate constants are determined from the slope and y-intercept respectively. Error ± SD. Polymerization conditions: 25 mM imidazole, pH 7.4, 50 mM KCl, 2.5 mM MgCl_2_, 1 mM EGTA, 2.5 mM MgATP,10 mM DTT, 0.25% methylcellulose, 2.5 mg/mL BSA, 0.5% Pluronic F-127, oxygen scavenging system (0.13 mg/mL glucose oxidase, 50 μg/mL catalase, and 3 mg/mL glucose), 37°C.

| | Critical concentration<br>$C_c$ ($\mu\text{M}$) | Disassembly<br>$k_-$ (subunits$\cdot\text{sec}^{-1}$) | Assembly<br>$k_+$ (subunits$\cdot\mu\text{M}^{-1}\cdot\text{s}^{-1}$) | n |
| --- | --- | --- | --- | --- |
| TgAct1 barbed (+) end rate constants |  |  |  |  |
| Prep 1 | 6.4 | 13.4 | 2.1 | 228 |
| Prep 2 | 7.8 | 17.4 | 2.2 | 373 |
| Prep 3 | 5.4 | 14.6 | 2.7 | 51 |
| Average | $6.5 \pm 1.2$ | $15.1 \pm 2.1$ | $2.3 \pm 0.3$ | |
| Skeletal actin barbed (+) end rate constants |  |  |  |  |
|  | 0.09 | 1.2 | 12.7 | 278 |
| TgAct1 pointed (-) end rate constants |  |  |  |  |
| Prep 1 | 84.3 | 10.8 | 0.1 | 111 |
| Prep 2 | 44.3 | 9.1 | 0.2 | 95 |
| Prep 3 | 91.9 | 11.9 | 0.1 | 43 |
| Average | $73.5 \pm 25.6$ | $10.6 \pm 1.4$ | $0.2 \pm 0.04$ | |

The rate constants and critical concentration of TgAct1 deviate significantly from those of skeletal muscle actin. For the TgAct1 barbed-end, the critical concentration is 6.5 ± 1.2 µM (**Fig. 1h, blue, Table 1**) which is ∼50-fold higher compared to skeletal muscle actin (29). We find that the disassembly rate constant at the barbed-end of TgAct1 filaments is 15.1 ± 2.1 sec^-1^ and the assembly rate constant (k^+^) is 2.3 ± 0.3 μM^-1^·s^-1^ which indicates that the barbed-end of TgAct1 disassembles about 15 times faster and assembles about 6 times slower compared to skeletal actin (29).

A linear fit to the pointed-end disassembly rate versus TgAct1 concentration shows that that the average critical concentration (x-intercept, 73.5 ± 25.6 µM) is ∼60-fold higher than the pointed-end of skeletal muscle actin, which arises from a disassembly rate constant (y-intercept, 10.6 ± 1.4 sec^-1^) that is 33-fold faster than skeletal actin (30) and a slow assembly rate constant (0.2 ± 0.04 sec^-1^) (**Fig. 1i, blue, Table 1**).

Given the high critical concentration, we measured the cellular concentrations of TgAct1 to assess if they are sufficient to support polymerization *in vivo*, in the absence of polymerization factors. As shown by western blot (**Fig. 2a**), *T. gondii* has a relatively high cytosolic actin concentration compared to other cell types (**Fig. 2b**), and this concentration would be sufficient to drive actin polymerization from both filament ends because the cellular concentration is above the critical concentration for polymerization at both filament ends.

### Rapid nucleotide turnover and exchange allows for intrinsic TgAct1 filament treadmilling

Next, we wished to understand how the ATPase properties of TgAct1 differ from skeletal actin to maintain filament treadmilling. We polymerized TgAct1 and monitored inorganic phosphate (Pi) release using an MESG assay (31, 32). For skeletal actin, the amount of phosphate released plateaus at an equimolar amount of F-actin (**Fig. 2c, orange**). This was expected because once F-actin formation plateaus, the rate of filament subunit disassembly, nucleotide exchange and reincorporation is negligible on this timescale. That is, polymerized subunits undergo a single ATPase turnover. Since nucleotide exchange does not occur in monomers incorporated into a filament, polymerized subunits generate inorganic phosphate once, and then must disassemble from the filament in order for nucleotide exchange to occur (33–35). Strikingly, TgAct1 phosphate release does not plateau, and each TgAct1 releases (on average) 3 phosphates over the 1-hour time course (**Fig. 2c, blue**). This indicates that each actin subunit hydrolyzes multiple ATP molecules during the timescale of the experiment, consistent with TgAct1 treadmilling. As a control, we measured the Pi release rate over a range of actin concentrations (**Fig. 2d**). As expected, the Pi release rate increases with actin concentration with steady-state ATPase rate (slope) of 8.5 x 10^-4^ μM^-1^·s^-1^.

MgADP-actin subunits that dissociate from filaments exchange bound MgADP for MgATP before incorporating to the barbed-end (30, 33, 35, 36). For skeletal actin, the rate constant for nucleotide exchange is slow (37–41) and generally requires exchange factors, such as profilin, for recharging the monomers with MgATP (42–44). Because TgAct1 rapidly treadmills in the absence of exchange factors (**Fig. 1e and Fig. 2c**), we hypothesized that nucleotide exchange from TgAct1 must be faster compared to skeletal actin. To test this hypothesis, ATP was exchanged for ethenoATP (MgεATP) which displays a fluorescence enhancement at 410 nm when bound to actin. The dissociation rate constant for εATP was determined by competing bound MgεATP a large molar excess of unlabeled MgATP and monitoring the change in fluorescence over time (40, 41, 45). The εATP dissociation from skeletal actin was slow (k_-εATP_ = 0.003 ± 0.0002 sec^-1^) (**Fig. 2e-f, orange**) consistent with previous results (33, 38). In contrast, the exchange of εATP from TgAct1 was over 50-times faster (k_-εATP_ = 0.16 ± 0.03 sec^-1^) (**Fig. 2e-f, blue**). The fast nucleotide exchange explains how TgAct1 can rapidly exchange bound nucleotide and become competent for filament assembly, even without nucleotide exchange factors.

### Optimization of experimental conditions for cryo-electron microscopy of Toxoplasma actin filaments

Next, we sought to use cryo-electron microscopy to determine the structural basis for the distinct polymerization properties of TgAct1. Initial negative stain micrographs in 0.1 mM ATP incubated for 1 hour at 37 °C with or without jasplakinolide reproduced what has been reported in the literature (22, 46); long filaments are visible with jasplakinolide, whereas short filaments assemble without jasplakinolide (**Ext. Data Fig. 3a and 3b**). Substituting the stabilizing, non-hydrolysable analog AMPPNP for ATP in the polymerization buffer produced filaments without jasplakinolide (**Ext. Data Fig. 3c**), but we were hesitant to use a non-native ligand for structural exploration because inclusion of 0.1 mM AMPPNP in the actin growth assay resulted in an ∼1.5-fold increase in the disassembly rate from the barbed-end and 2-fold increase in the critical concentration compared to MgATP conditions (**Ext. Data Fig. 4**).

**Figure 3:**
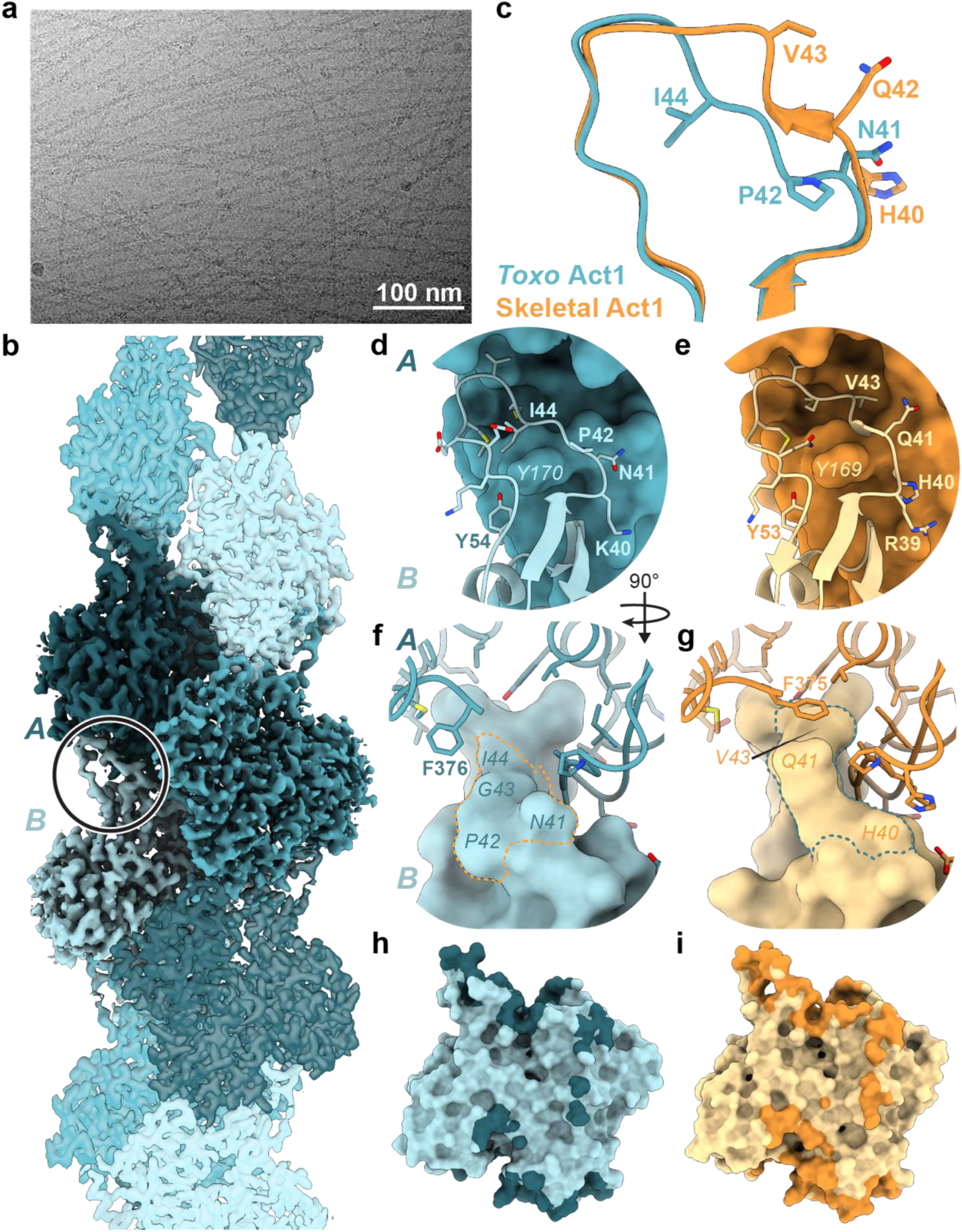
Comparison of unstabilized TgAct1 filament to skeletal actin filaments. **a**, Micrograph of TgAct1 filaments in the presence of 1 mM Mg-ATP. **b,** Reconstruction of TgAct1 filaments in the presence of 1 mM Mg-ATP; protomers in shades of blue. The circle indicates the location of the D-loop in one protomer. **c,** Overlay of D-loops from unstabilized TgAct1 filament model (blue) and the skeletal actin filament 8d13 (orange). **d/e,** View of the D-loop (ribbon and sticks) and binding pocket (surface) from TgAct1 (d, blue) and skeletal actin (e, orange). D-loop residues within 5 Å of the pocket are shown as sticks. **f/g,** View of the D-loop (surface) and binding pocket (ribbon and sticks) from TgAct1 (f, blue) and skeletal actin (g, orange). Residues within 5 Å of the D-loop are shown as sticks, except for TgAct1 F376, which is shown for illustrative purposes only. Dotted lines indicate the locations of D-loop residues 41-44 (TgAct1) and 40-43 (skeletal actin). **h/i,** Surface representation of a protomer of TgAct1 (h, blue) and skeletal actin (i, orange), with the shaded region showing the buried surface area for each protomer.

**Figure 4:**
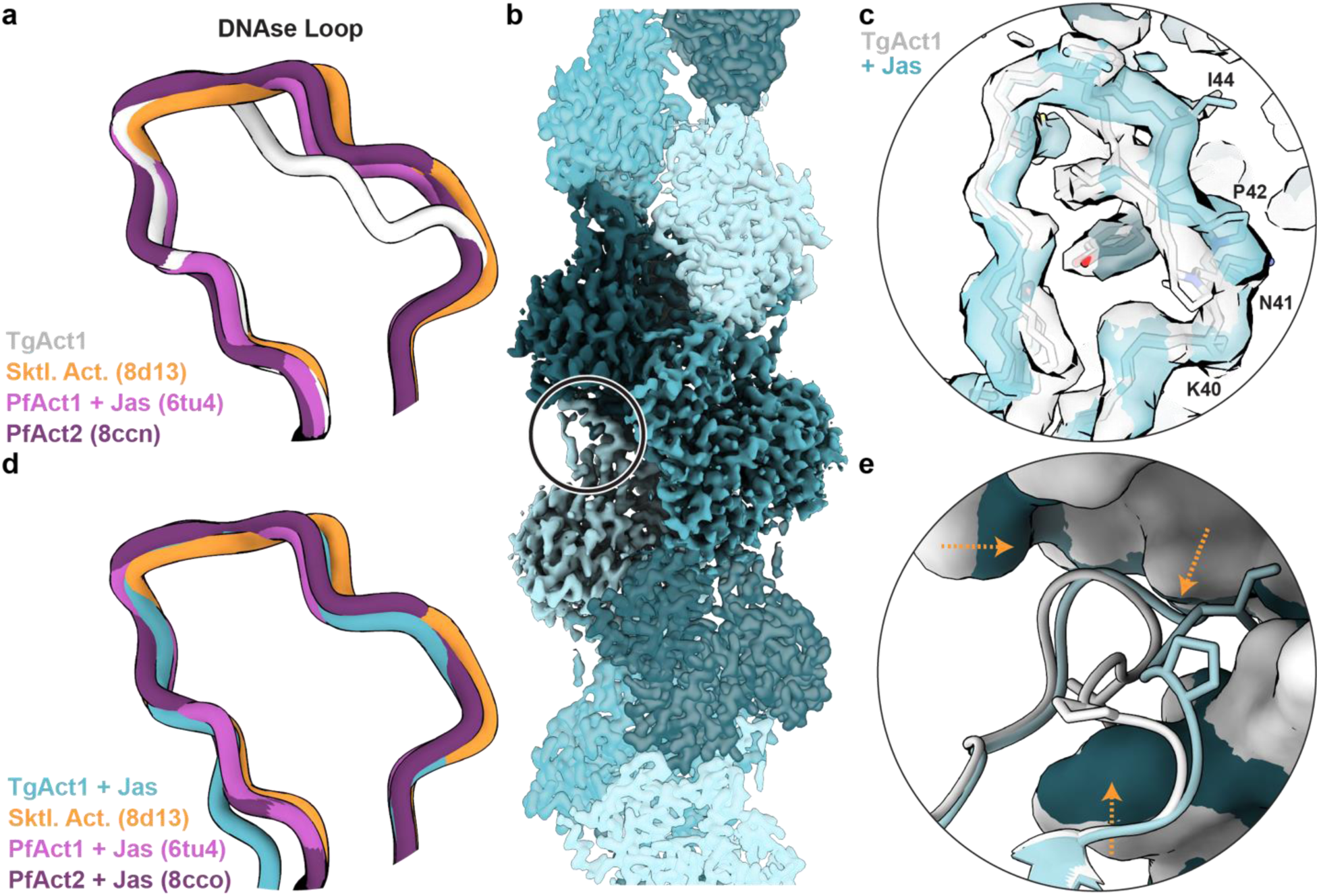
Comparison of stabilized and unstabilized TgAct1 filaments to actin filaments from other species. **a**, The D-loops from unstabilized TgAct1 (gray), chicken skeletal actin (orange, PDB ID 8d13), *P. falciparum* Act1 + jasplakinolide (magenta, PDB ID 6tu4), and *P. falciparum* Act2 (purple, PDB ID 8ccn). **b,** Reconstruction of TgAct1 filaments in the presence of 33 µM jasplakinolide and 0.1 mM Mg-ATP; protomers in shades of blue. The circle indicates the location of the D-loop in one protomer. **c,** Overlay of volume from the D-loops of TgAct1 + jasplakinolide (blue) and unstabilized TgAct1 (gray); C, CA, and N backbone atoms from residues 36-55 in each model shown and selected residues shown as sticks. **d,** The D-loops from TgAct1 + jasplakinolide (blue), chicken skeletal actin (orange, PDB ID 8d13), *P. falciparum* Act1 + jasplakinolide (magenta, PDB ID 6tu4), and *P. falciparum* Act2 + jasplakinolide (purple, PDB ID 8cco). **e,** Overlay of the D-loop (ribbon) and binding pocket (surface) from unstabilized TgAct1 filaments (gray) and TgAct1 + jasplakinolide filaments (blue). Pro42 and Ile44 shown as sticks. Orange arrows indicate the direction of motion.

Based on our kinetic data, we tested filament conditions that might slow filament disassembly. Shortening the incubation period and lowering the assembly temperature to 25°C did not produce any filaments (**Ext. Data Fig. 3d**). Decreasing the incubation time and increasing the ATP concentration produced promising results. At time points between 10 and 30 minutes, long filaments are observed in negative stain, with some filaments extending several microns (**Ext. Data Fig. 3e and 3f**). This observation suggests that due to the treadmilling, ATP hydrolysis and nucleotide exchange are rapid enough for TgAct1 to consume the majority of the 0.1 mM ATP in 1 hour in the initial reaction conditions. Given the steady-state ATPase rate (**Fig. 2d**, 8.5 x 10^-4^ μM^-1^·s^-1^), 33 µm TgAct1 should hydrolyze 0.1 mM MgATP in one hour. Decreasing the incubation time and increasing the ATP concentration allowed us to circumvent this outcome. Thus, to capture long filaments in cryo conditions, we froze TgAct1 samples after a 10 to 20-minute incubation period with 1 mM ATP (**Fig. 3a**).

### Structure of unstabilized ADP-bound TgAct1 filaments reveal altered D-loop positioning and decreased buried surface area

We determined the structure of unstabilized TgAct1 filaments at 2.5 Å resolution, which revealed an overall architecture similar to skeletal and other actin filament structures (**Fig. 3b**, **Ext. Data Fig. 5 and 6A**, **Table 2**). TgAct1 and skeletal muscle actin share 83% sequence identity (4), and the structures show many similarities. As we identified MgADP in the nucleotide binding site (**Ext. Data Fig. 6b**), we compared TgAct1 to structures of rabbit and chicken skeletal muscle actin bound to MgADP (PDB IDs 8d13 and 8a2t) (47, 48). A single TgAct1 protomer is in nearly the same conformation as skeletal actin, with Cα RMSD values of 0.5 and 0.8 for chicken and rabbit actin, respectively (**Supplemental Movie 3**). The helical symmetry of TgAct1, 28.1 Å rise and –167.1° twist, is nearly identical to chicken and rabbit MgADP actin filaments, at 28.1 and 27.5 Å rise, respectively, and –166.7° twist for both.

**Figure 5:**
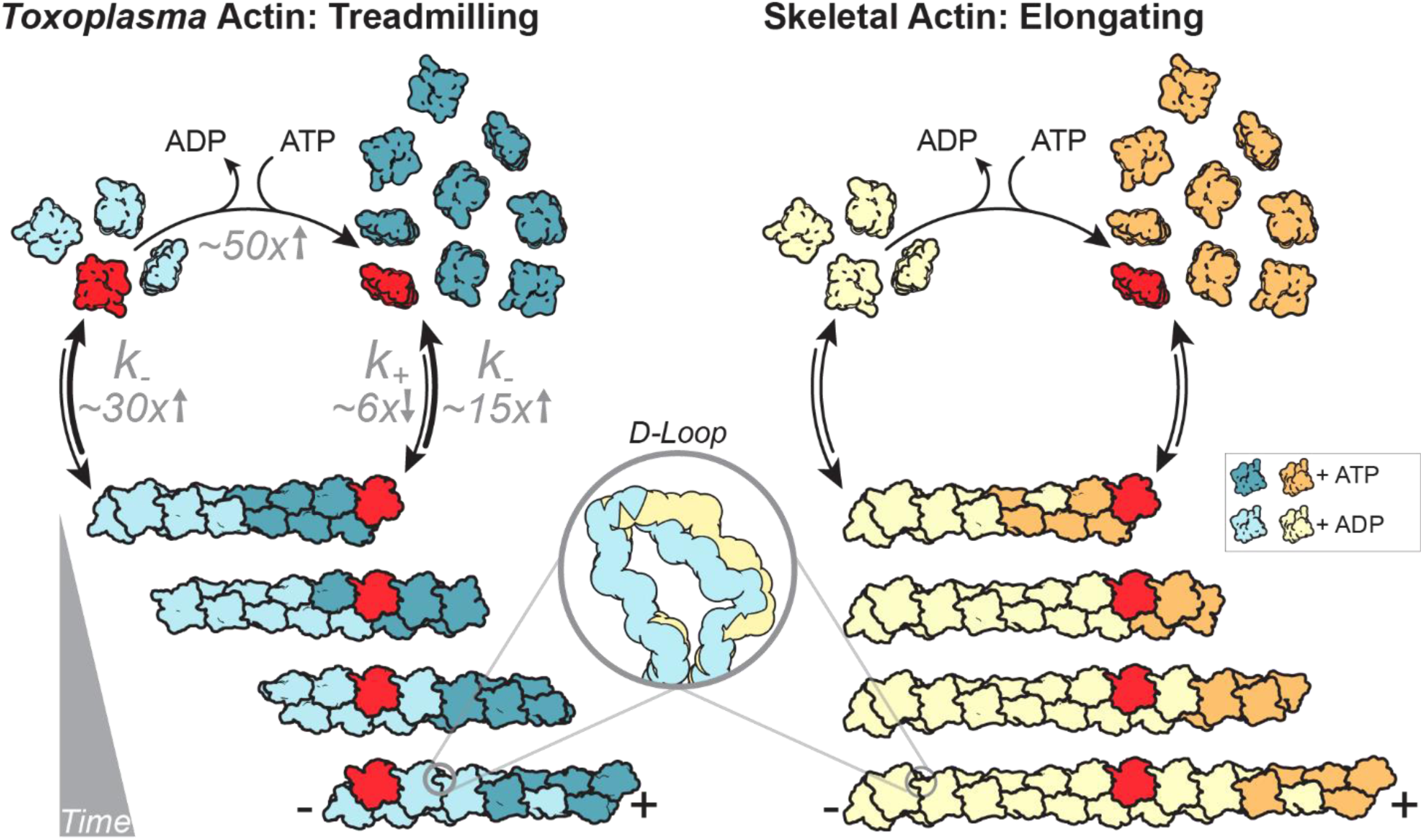
The Filament Properties of Toxoplasma Actin versus Skeletal Actin. A low barbed-end assembly rate and a high pointed-end disassembly rate leads to a high critical concentration for TgAct1 relative to skeletal actin – an effect mediated in part by changes within the D-loop of TgAct1. Paired with the ability to rapidly exchange nucleotide, actin filaments in *T. gondii* rapidly treadmill at concentrations where skeletal actin filaments elongate. Change in rates of TgAct1 assembly, disassembly and nucleotide exchange compared to skeletal muscle actin are indicated in grey text.

**Table 2:**
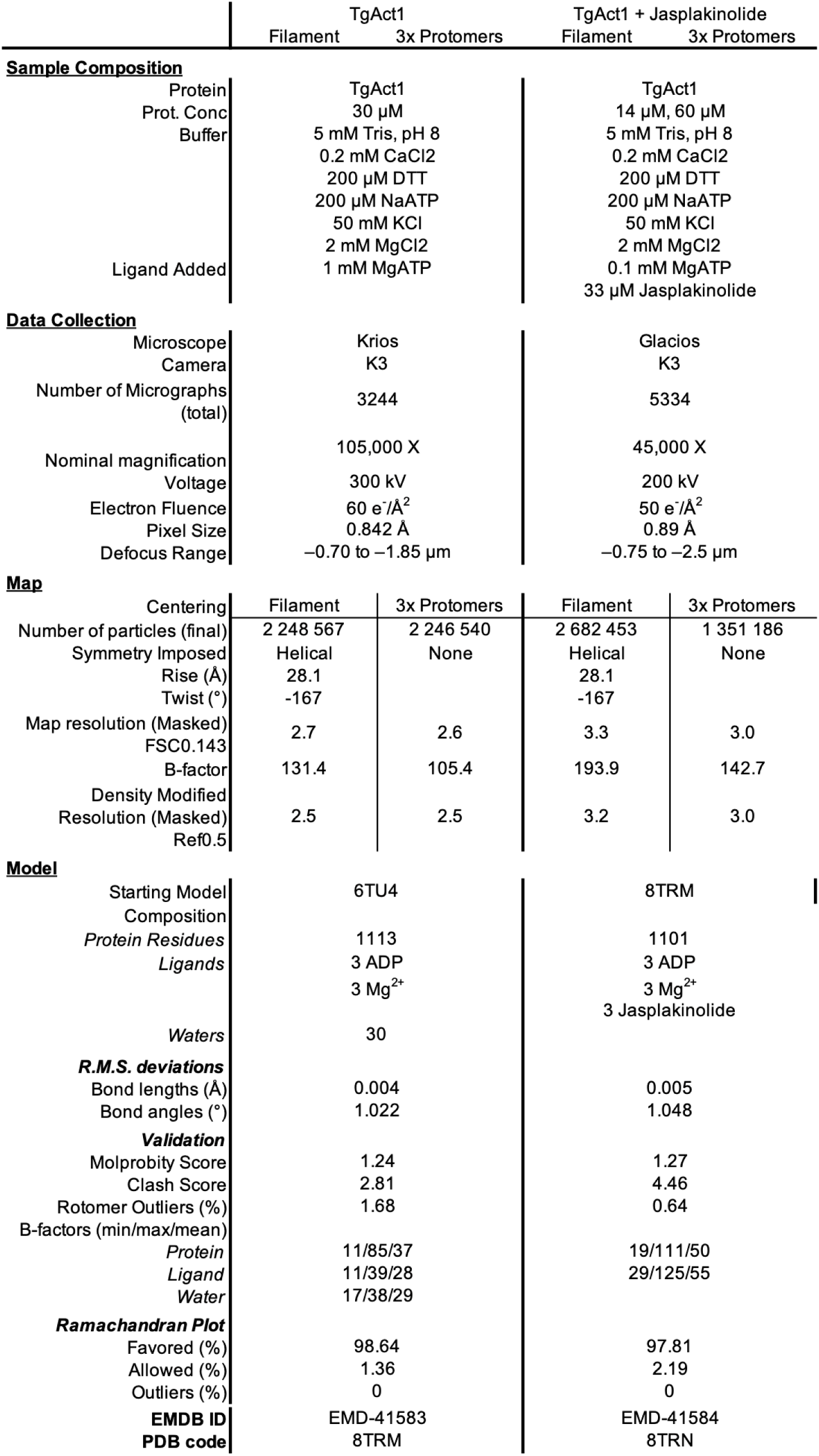
Cryo-EM data collection, refinement, and validation statistics.

However, there are notable differences between the structures. The most striking difference is in the position of the D-loop (residues 40-51 in TgAct1 and residues 39-50 in skeletal muscle actin), which forms important longitudinal contacts between actin protomers (**Ext. Data Fig. 6c-d**). In TgAct1 filaments, the D-loop has shifted, moving the loop partially out of the pocket where the loop is typically bound, with a Cα RMSD value of 2.1 when compared to chicken skeletal actin (47). There are six amino acid substitutions on the D-loop in TgAct1 relative to skeletal actin (**Ext. Data Fig. 7**). Two of these substitutions, amino acids Pro42 and Ile44, point in the opposite direction relative to the comparable residues in skeletal actin (Gln41 and Val43) (**Fig. 3c**). While Val43 of skeletal actin sits within a hydrophobic portion of the binding pocket, the comparable residue in TgAct1 (Ile44) is displaced by 5 Å and does not make inter-protomer contacts (**Fig. 3d-e**). These changes in the D-loop conformation break most of the backbone interactions that occur among skeletal actin D-loop residues 41-44 (TgAct1 42-44), and hydrogen bond contacts between side chains are lost due to a combination of residue substitution and mainchain displacement (TgAct1 amino acids 39-41 and 46) (**Fig. 3f-g, Ext. Data Fig. 8a and b**). In contrast to the substitutions on the D-loop, the amino acids in the pocket of the longitudinally adjacent protomer where the D-loop binds are identical between TgAct1 and skeletal actin (**Fig. 3f-g**). Overall, this reduces the buried surface area in this interaction by about one third, from 638 Å^2^ in chicken skeletal actin (PDB ID 8d13) to 485 Å^2^ in TgAct1. The D-loop position and resulting decrease in longitudinal protomer contacts likely contribute to the decreased filament stability of TgAct1, explaining the rapid depolymerization rate constant and high critical concentration.

Other structural differences between TgAct1 and skeletal actin filaments are more modest. The C-terminal phenylalanine of TgAct1 occupies a different position than skeletal actin and in TgAct1 it does not interact with the D-loop as it does in skeletal actin (**Fig. 3f-g**). Tyr54 from TgAct1 (skeletal actin Tyr53) establishes a hydrogen bond with Tyr170 (skeletal actin Tyr169) on the neighboring protomer (**Ext. Data Fig. 8a and b**). There are four substitutions on the hydrophobic loop (TgAct1 residues 263-275), but the mainchain retains the same conformation. The substituted side chains only make intraprotomer contacts; however, one of the substitutions (TgAct1 Lys270 – skeletal actin Met 269) adds a hydrogen bond with the backbone of Asp180, subtly shifting the mainchain around residues 177-180 (**Ext. Data Fig. 8c**). Comparing the total buried surface area for one protomer of TgAct1 versus one protomer of skeletal actin provides a summation of the changes: TgAct1 loses 364 Å^2^ of buried surface area relative to skeletal actin filaments (8d13: 3115 Å^2^; TgAct1: 2751 Å^2^), the majority of which is from the change in the position of the D-loop (**Figure 3h-i**).

### The D-loop of Jasplakinolide-stabilized TgAct1 filaments closely resembles skeletal and P. falciparum

TgAct1 has 93% sequence identity to *Plasmodium falciparum* actin 1, and the D-loops and the pocket where the D-loops bind are identical. (**Ext. Data Fig. 7**). Despite this identity, the D-loops of native TgAct1 and *P. falciparum* Act1 stabilized with jasplakinolide have different conformations (**Fig. 4a, Ext. Data Fig. 8d**). The D-loop conformation of jasplakinolide-stabilized *P. falciparum* Act1 is similar to the D-loops of both unstabilized skeletal actin filaments and the second *P. falciparum* actin isoform, Act2. Skeletal actin has a distinct D-loop conformation when bound to jasplakinolide (**Ext. Data Fig. 8e**). This suggests that the effect of jasplakinolide on actin filaments are not uniform across actin isoforms and that its effects on TgAct1 may differ from skeletal or *P. falciparum* actins. To test this hypothesis, we solved a structure of TgAct1 bound to jasplakinolide at 3.0 Å (**Fig. 4b, Ext. Data Fig. 6, 6a-f, Table 2**). Jasplakinolide-bound filaments retained the same helical parameters as unstabilized filaments (28.1 Å rise and –167° twist). The nucleotide binding site contained MgADP (**Ext. Data Fig. 6b**) and the protomers of the filaments are very similar, with a Cα RMSD of 0.5 Å. As expected, jasplakinolide binds in the same location in TgAct1 filaments as it is found in skeletal and *P. falciparum* filaments (**Ext. Data Fig. 6f**). It primarily bridges the lateral interaction between protomers and increases the lateral surface contacts from 702 Å^2^ in unstabilized filaments to 1016 Å^2^ in jasplakinolide-bound filaments.

The D-loop of the jasplakinolide-bound filament adopts a conformation nearly identical to that of jasplakinolide-bound *P. falciparum* Act1 and skeletal actin loops that is distinct from the unstabilized TgAct1 loop (**Fig. 4c and d, Ext. Data Fig. 6c-d**). Cα RMSD values comparing the D-loops demonstrate the overlap: 0.8 and 1.2 comparing jasplakinolide-bound TgAct1 to jasplakinolide-bound *P. falciparum* Act1 and native chicken skeletal actin, respectively. With jasplakinolide bound, amino acids Pro42 and Ile44 point into the binding pocket, though the volume does suggest some mixed occupancy with the unstabilized D-loop conformation that we were unable to resolve by classification (**Ext. Data Fig. 6c-d**). There is also a shift in residues 51-53 outside of the D-loop, which is not seen in other structures. Here, the volume suggests that the Cys53 may have become oxidized, possibly leading to the change (**Ext. Data Fig. 6g**). The C-terminus of the protein is unresolved past residue His372 in the jasplakinolide-bound structure (**Ext. Data Fig. 6h**). More subtle shifts occur among the subdomains within one protomer. The overall effect is for residues 139-144 and 169-173 in subdomain 2 and residues 352-370 in subdomain 1 to pinch the D-loop in its binding pocket, likely stabilizing its position (**Fig. 4e**), which suggests that the TgAct1 D-loop can adopt at least two conformations within the filament.

Neither actin isoform from *P. falciparum* shows large differences in the positioning of the D-loop when compared to skeletal actin filaments (24, 49, 50). For *P. falciparum* Act2, the substitution of an asparagine for the glycine at position 42 might facilitate a more stable filament. In contrast, biochemical work on *P. falciparum* Act1 suggests that the D-loop contributes to the instability of the filaments (21, 24). Because of the high identity of *P. falciparum* Act1 to TgAct1, it appears possible that in unstabilized filaments the *P. falciparum* Act1 D-loop could adopt the same position as the unstabilized TgAct1 D-loop.

## Discussion

### Properties of TgAct1 filaments

In this study, we show that TgAct1 is capable of polymerizing into long filaments in the absence of assembly factors, stabilizers, or the actin chromobody, which is currently the only actin probe capable of detecting actin filaments in Apicomplexan parasites. Unlike skeletal actin, TgAct1 filaments rapidly treadmill at concentrations of actin above 12 µM (**Fig. 5**). This unusual feature has also been observed for *P. falciparum* Act1 (21) which shares 93% sequence identity to TgAct1 and thus, appears to be a conserved feature of Apicomplexican actin.

The structural basis for rapid disassembly is likely explained by the conformation and dynamics of the D-loop. Our structures reveal two distinct D-loop conformations for the stabilized vs. unstabilized structure: in the jasplakinolide stabilized structure, the D-loop is fully inserted into the binding pocket, whereas in the absence of stabilizing agents, the loop is partially out of the binding pocket. It suggests that the loop makes the full set of contacts during filament assembly and shifts after assembly, decreasing the number of longitudinal interactions. Given the role of the D-loop in regulating F-actin stability through longitudinal contacts, we propose that these structural differences may account for the reduced filament stability and high dissociation constants at the pointed-end that drive treadmilling (**Fig. 5, inset**). Further support for this idea comes from data showing that replacing the *P. falciparum* Act1 D-loop with the mammalian sequence has a stabilizing effect on the filament (24) accompanied by a ∼5-fold reduction in the critical concentration (21). Despite the known importance of the D-loop in regulating *P. falciparum* Act1 filament stability, previous structures of *P. falciparum* Act1 stabilized with jasplakinolide observed minimal differences in the position of the D-loop of *P. falciparum* Act1 relative to skeletal muscle actin (49). The D-loop differences between stabilized and unstabilized TgAct1 filaments helps to reconcile why previous structures of *P. falciparum* Act1 could not fully explain the importance of the D-loop in filament stability.

In order to support rapid treadmilling, TgAct1 monomers must quickly release ADP and recharge with ATP in the absence of nucleotide exchange factors. Exploration of the hinge regions involved in the F to G transition did not reveal any substantive differences between TgAct1 and skeletal actin. There is only one substitution within the nucleotide binding pocket between skeletal muscle actin and TgAct1: a leucine (Leu16) is substituted with an asparagine (Asn17) in TgAct1 (**Ext. Data Fig. 7 and 8f**). *P. falciparum* Act1, like TgAct1, also contains this substitution (24). Comparison of the gelsolin-bound actin structures of *P. falciparum* Act1 and skeletal actin (51, 52) to the stabilized filament structure of *P. falciparum* Act1 and the unstabilized filament structure of TgAct1 suggests how the residue might affect the nucleotide exchange rate. The G-actin structure of *P. falciparum* Act1 shows Asn17 hydrogen bonding with the backbone carbonyl of Phe34 in the adjacent beta strand. For this change to occur, the hydrogen bond with the alpha phosphate in the filament protomer must be broken and the gamma carbon of Asn17 must move 1.6 Å toward the backbone carbonyl of Phe34. In contrast, the gamma carbon of hydrophobic Leu16 only moves half that distance (**Ext. Data Fig. 8g**). The substitution of the asparagine for the leucine and the associated shifts in G-actin unbury part of the nucleotide, and decrease the hydrophobic character while widening a portion of the binding pocket, potentially facilitating nucleotide exchange.

### Conflicting views on the properties of Apicomplexan actin

This work challenges the idea that TgAct1 forms only short unstable filaments. We consistently observe filaments longer than 20 µm using fluorescent microscopy, and filaments longer than 1 µm that span the micrographs in negative stain electron microscopy. In addition, bulk ATPase assays indicate that 24 µM TgAct1 monomers treadmill into and out of the filament three times per hour. Given the dissociation constant from the pointed-end (11 sec^-1^), the average filament length under these conditions is estimated to be ∼36 µm. Previous methods for measuring TgAct1 filament length were either indirect, using centrifugation, or by negative stain electron microscopy of filaments near their critical concentration, where low availability of monomers limit their length (46). It is unclear whether the N-terminal HIS tag used in previous studies influences TgAct1 or *P. falciparum* Act1 filament stability and length.

While there are differences in the reported critical concentration for Apicomplexan actins, our results are largely consistent with the high critical concentration obtained by Lu et al. for *P. falciparum* Act1 (21). Previous experiments that found low critical concentrations for TgAct1 (22) or *P. falciparum* Act1 (23) were performed over time courses many hours in length. Our ATPase results indicate that unlike skeletal actin, TgAct1 will be completely converted to MgADP (**Fig. 2c**) and long filaments will be lost over these long time periods because ADP-actin has a 10-fold increased critical concentration compared to ATP-actin (21, 30). In addition, monitoring polymerization of apicomplexican actin overnight is also problematic because small oligomers form when not stabilized by the addition of ammonium acetate (23), which can complicate the interpretation of light scattering or pyrene fluorescence.

### Implications for rapidly treadmilling filaments in T. gondii parasites

*In vivo* visualization of actin in *T. gondii* using the actin chromobody has revealed a vast network of highly dynamic actin that is well supported by its *in vitro* properties (12). Because the concentration of TgAct1 in the cell is near the critical concentration for the pointed-end, its assembly and disassembly may be locally fine-tuned by actin sequestering proteins which could function to drive the dynamics of F-actin in *T. gondii*. Apicomplexan parasites have an unusually limited repertoire of actin binding proteins (28). *T. gondii* encodes a single gene of ADF and profilin, and three formins. In mammalian cells, profilin binds to G-actin and plays a dual function in nucleotide exchange and enhancement of formin-mediated actin polymerization (53–55). Considering our finding that TgAct1 has an extremely fast nucleotide exchange rate, the function of profilin may be restricted to enhancing actin polymerization from one of the three formins in the cell.

The actin-depolymerizing factor (ADF)/cofilin is a family of proteins that regulate the turnover of actin networks in the cell (56–59). ADF/cofilin proteins typically bind F-actin and destabilize filament networks by severing actin filaments and accelerating depolymerization from the pointed-end. ADF is essential to the survival of *T. gondii* and its depletion promotes the assembly of F-actin structures in the cell (7, 16). Interestingly, Apicomplexican ADF lacks the residues for binding F-actin and therefore shows very weak severing activities. Instead, it exclusively binds and sequesters G-actin (7, 60). Severing activities was likely lost because of the high disassembly rates of TgAct1, which did not necessitate severing for fast disassembly. Thus, ADF functions primarily in *T. gondii* to regulate the pool of available actin monomers.

A key property of TgAct1 that makes it primed for rapid disassembly is the ∼30-fold higher disassembly rate for the pointed-end compared to skeletal actin. Thus, below the critical concentration, TgAct1 filaments will depolymerize ∼30-times faster than conventional actin. Considering our findings that the cellular concentration of TgAct1 is less than 2-fold from the critical concentration of pointed-end, small changes in the expression of ADF, or its activation through post-translational modification may allow for rapid transitioning from filament to monomer without the need for actin severing proteins or disassembly factors.

Why does Apicomplexican actin have an unusually high critical concentration? This novel property would seemingly require the expression of roughly 10-times more actin and regulatory proteins to generate filaments which is energetically wasteful. We propose that the high critical concentration is a functional consequence of the high monomer disassembly rates. While more actin expression is required for filament growth, the fast disassembly rates allow the parasite to quickly disassemble the actin network with a minimal set of actin regulatory factors. Such a mechanism could have evolved due to constraints of the lytic cycle on parasite survival. Intracellular parasites contain an extensive and dynamic actin network. When calcium ionophore is used to trigger parasite egress the network rapidly disassembles in less than 60 seconds, prior to the onset of parasite motility (16). During motility and egress, actin is enriched at the parasites basal end (18). Thus, parasite motility and the transition from the intracellular to extracellular states (and vice versa), necessitate fast reorganization of the actin network and transitions between F and G-actin.

Due to its ubiquity across the domains of life, actin serves as a model for evolutionary tuning of the dynamic properties of a conserved polymer. In general, eukaryotic actin is highly conserved at the sequence level and in the overall structure of the filament, and achieves functional diversity in part through interactions with a host of regulatory proteins that tune its dynamic properties. Bacterial actins, on the other hand, typically have many fewer interacting partners, which has allowed evolutionary diversification to yield different dynamic properties through sequence changes that introduce significant changes to the filament architecture (61). TgAct1 sits between these two extremes, demonstrating that smaller evolutionary changes can give rise to new dynamic properties without large changes to the overall filament structure.

## Methods

### DNA expression constructs

TgAct1 (ToxoDB: TgGME49_209030) was fused as its C-terminus to a 14 amino acid GS linker sequence followed by a β-thymosin–6xHIS tag and inserted into pFastBac (Thermo Fisher Scientific) behind the p10 promoter for expression in Sf9 cells. The β-thymosin fusion strategy was previously used for the expression of *Plasmodium falciparum* and *Dictyostelium* actin, which shares 95% and 85% identity to TgAct1 respectively (21, 25, 26). The Actin-chromobody (ChromoTek Inc., Hauppauge, NY) containing a C-terminal EmeraldFP–6xHIS tag was cloned into pET22b (Novagen) for expression in *E. coli* BL21(DE3) (16).

### Protein expression and purification

Expression and purification of untagged TgAct1 was adapted from previous methods (21, 25). Two billion Sf9 cells were infected with virus and grown for 3 days at 27°C. Cells were harvested by centrifugation at 2000 RPM and resuspended in 50 mL lysis buffer (10 mM HEPES, pH 8.0, 0.25 mM CaCl_2_, 0.3 M NaCl, 0.5 mM DTT, 0.5 mM Na_2_ATP, 5 µg/mL leupeptin, 1x cOmplete ULTRA Tablets (Roche)). Cells were lysed by sonication in an ice water bath and then clarified at 250,000 x g for 35 minutes. The supernatant was then added to 3 mL of HIS Select Nickel affinity gel (Sigma) equilibrated in wash buffer (10 mM HEPES, pH 8.0, 0.25 mM CaCl_2_, 0.3 M NaCl, 0.5 mM DTT, 0.25 mM Na_2_ATP) for 45 minutes at 4°C. The affinity gel was then lightly sedimented at 4°C, 600 RPM and added to a column and washed with 15 bed volumes of wash buffer followed by 6 bed volumes of wash buffer containing 10 mM Imidazole (pH 8). The protein was eluted with wash buffer containing 200 mM Imidazole, pH 8. Peak fractions were concentrated to 2 mL Amicon Ultra-50 (Millipore) and dialyzed against 1 L G buffer overnight (5 mM Tris, pH 8.2, 0.25 mM CaCl_2_, 0.5 mM DTT, 0.25 mM Na_2_ATP, 1 µg/mL leupeptin, 4°C). Protein was divided into 150 µL aliquots, snap frozen in liquid nitrogen and stored at –80°C. Small scale chymotrypsin digests were performed to determine the optimal molar ratios of chymotrypsin:TgAct1–β-thymosin–HIS for removal of the β-thymosin–HIS tag. Volumes of 20 µL of actin were incubated with chymotrypsin (Sigma) at the indicated molar ratio (**Extended Data** Fig. 1a) in a 27°C water bath for 15 minutes and then quenched with 0.5 mM PMSF (Roche). Samples were analyzed on a Coomassie stained SDS-PAGE PAGE gel. Once an optimal molar ratio was determined, the remaining prep was digested, quenched with PMSF, clarified 400,000 x g for 30 minutes and applied to a Resource Q (GE Healthcare) ion exchange column equilibrated with G buffer. TgAct1 was separated from the cleaved tag by applying a 0–0.5 M NaCl salt gradient in G buffer. Fractions containing purified tagless TgAct1 were concentrated to 0.5 mL and dialysed overnight against G-buffer containing 0.2 M ammonium acetate, pH 8, 4°C, which prevents spontaneous actin oligomerization (21, 23). The following day, the sample was clarified 400,000 x g for 30 minutes and applied to a Superdex 200 Increase 10/300 GL (GE Healthcare) gel filtration column equilibrated in G-buffer containing 0.2 M ammonium acetate. Fractions corresponding to monomeric actin were pooled, concentrated in an Amicon Ultra – 15 (Millipore), snap frozen in liquid nitrogen and stored at –80°C.

The actin chromobody was purified from E. coli, BL21(DE3). Transfected cells were grown to mid log (OD = 0.7) and induced with 0.4 mM IPTG (Roche) and grown overnight at 16°C. Cells were harvested by centrifugation at 3500 RPM and resuspended in 40 mL lysis buffer (10 mM NaPO_4_, 300 mM NaCl, 0.5% glycerol, 7% sucrose, pH 7.4 at 22°C, 0.5% NP40, 0.5 mM PMSF, 1x protease inhibitor tablets (Pierce)). Lysozyme (Sigma) was added to 3 mg/mL and incubated at 4°C for 30 minutes. The cell lysate was then sonicated in an ice water bath and then clarified at 250,000 x g for 35 minutes. The supernatant was then added to 3 mL of HIS Select Nickel affinity gel (Sigma) equilibrated in wash buffer (10 mM NaPO_4_, 300 mM NaCl, 0.5% glycerol, pH 7.4 at 22°C, 0.5 mM DTT) for 45 minutes at 4°C. The affinity gel was passed over a column and washed with 20 bed volumes of wash buffer followed by 10 bed volumes of wash buffer containing 10 mM Imidazole (pH 7.4). The protein was eluted with wash buffer containing 200 mM Imidazole, pH 7.4. Peak fractions were concentrated to 3 mL and dialyzed against 0.2 L glycerol storage buffer (25 mM Imidazole, pH 7.4, 300 mM NaCl, 50% glycerol, 1 mM DTT). The final protein concentration was determined using a Bradford reagent (Thermo Scientific).

### *In vitro* actin growth assay

TgAct1 was thawed and clarified at 400,000 x g for 30 minutes. The storage buffer was exchanged with G buffer using a Zeba spin-desalting column (Thermo Fisher) and the actin was stored on ice for up to 4 hours. The concentration of the actin was determined by absorbance at 280 nm (1 OD = 38.5 µM). It should be noted that determining actin concentration using Bradford gives an measurement approximately 2-fold less than that measured by absorbance which could account in part for ∼2-fold difference in the calculated *C_c_* between Plasmodium actin (21) and *Toxoplasma* actin (this study). The visualization of actin growth *in vitro* was adapted from previous studies on Plasmodium actin (21, 25). Flow chambers were bound to minimal densities of NEM treated muscle myosin, which is ATP-insensitive and functions to keep filaments in the imaging plane. The flow cells were then rinsed with ice cold buffer B (25 mM Imidazole, pH 7.4, 50 mM KCl, 2.5 mM MgCl_2_, 1 mM EGTA, 10 mM DTT), and blocked with buffer B containing 5 mg/mL BSA and 1% Pluronic F127. The indicated concentrations of G actin diluted in G buffer was mixed 1:1 with 2x polymerization buffer (50 mM Imidazole, pH 7.4, 5% methylcellulose, 100 mM KCl, 5 mM MgCl_2_, 2 mM EGTA, 20 mM DTT, 5 mg/mL BSA, 1% Pluronic F127, 5 mM MgATP, 50 nM actin chromobody-EmeraldFP, and an oxygen scavenging system (0.13 mg/mL glucose oxidase, 50 μg/mL catalase, and 3 mg/mL glucose), and passed twice through the flow chamber which was then immediately sealed with nail polish. Actin polymerization which was visualized by low concentrations of actin chromobody was imaged at 37°C in epifluorescence using a DeltaVision Elite microscope (Cytiva) built on an Olympus base with a 100x 1.39 NA objective and definite focus system. Images were acquired every 5 seconds for 5 to 30 minutes with a scientific CMOS camera and DV Insight solid state illumination module.

### Epifluorescence microscopy data processing

Movies of actin growth were analyzed using the ImageJ plugin MTrackJ (62). The rate of shrinkage or growth for each filament was determined by measuring the change in filament length at each end over the length of the 5-minute movie. The length of TgAct1 or skeletal actin was converted to number of actin subunits in Excel with 2.76 or 2.74 nm length increase per subunit respectively (24). Pauses in depolymerization at the pointed-end were excluded from rate determinations. Individual measurements were imported into GraphPad Prism (GraphPad Software ver. 10.0.0, Boston, Massachusetts USA) for statistical analysis and graphing. Kymographs of actin dynamics were generated using the imageJ plugin multiplekymograph.

### Measuring the rate of dissociation of ethenoATP from actin

For the preparation of ethenoATP (εATP) bound actin, TgAct1 or skeletal actin was clarified at 400,000 x g for 30 minutes and exchanged into G buffer as described above. Actin was diluted to 20 µM in 200 µL and calcium was exchanged with magnesium by simultaneously mixing in 0.2 mM EGTA and 80 µM MgCl_2_ followed by incubating on ice for 5 minutes. Free ATP was depleted from solution by the addition of 10 µL DOWEX resin that was washed and equilibrated in a solution of 2 mM Tris pH 8.0. The mixture was continuously inverted for 2 minutes and spun 12,000 RPM for 1 minute at room temperature to remove the resin. To the supernatant, 50 µM εATP (Jena Bioscience) was added and the mixture was incubated on ice for 30 minutes. For measuring nucleotide exchange, 150 µL εATP actin was added to a 96 well plate and 50 µL of 2 mM MgATP was added using an auto injector in a SpectraMax® i3x plate reader. The plate was then simultaneously mixed and measured for fluorescence at 410 nm resulting in a delay time of 0.2 seconds after injection. Fluorescence readings were taken at 0.2 second intervals for 5 minutes. Time courses were fitted to single exponentials, yielding the nucleotide dissociation rate constant from the observed rate constant of the best fit.

### Actin Pi release assay

TgAct1 or skeletal actin was clarified at 400,000 x g for 30 minutes and exchanged into fresh G buffer as described above. The actin solution was warmed for 30 seconds in a 37°C water bath and induced for polymerization by adding 10x polymerization buffer (250 mM Imidazole, pH 7.4, 500 mM KCl, 20 mM MgCl_2_, 10 mM EGTA, 2 mM MgATP, 10 mM DTT) to a 1x final concentration at 37°C. The amount of Pi released from actin was measured by absorbance at 360 nm every 20 seconds for 30 minutes to 1 hour using an EnzChek phosphate assay kit (Thermo Fisher) (63, 64) in a GENESYS™ 10 UV-Vis Spectrophotometer (Thermo Scientific) maintained at 37°C.

### Western blotting to determine the cellular actin concentration

Cells were syringe released, counted, and harvested by centrifugation at 3,500 r.p.m. for 4 minutes in a benchtop centrifuge. Cells were washed in PBS, and boiled for 7 minutes in SDS-PAGE loading buffer. Cell lysates equivalent to 2.5 x 10^6^ and 5 x 10^6^ cells/lane were run on SDS-PAGE next to known amounts of purified TgAct1 ranging from 50 to 300 ng as determined by OD280. Protein was then transferred to a nitrocellulose membrane, blocked with blocking buffer (2.5% non-fat milk in PBS) and incubated with a 1/1000 dilution in blocking buffer of rabbit anti-actin-antibody (Gift from Marcus Meissner) which binds TgAct1 (8). The blot was then washed, incubated with a 1/3000 dilution of goat anti-rabbit secondary antibody in blocking buffer conjugated to IRDye® 680RD (LI-COR), washed, and imaged on a Typhoon imager (Cytiva). Densitometry analysis was performed using imageJ to generate a standard curve of intensities from known actin concentrations and the ng amount of actin per lane was determined by a standard fit to the curve. Number of parasites per lane and cell volume determined previously (66) was used to calculate the cellular actin concentration.

### Preparation of rhodamine-labeled TgAct1 filaments

TgAct1 was thawed and desalted twice into a storage buffer (5 mM Tris, pH 8.2, 0.2 M ammonium acetate, 0.25 mM CaCl_2_, 0.5 mM DTT, 0.25 mM Na_2_ATP, 4°C) lacking DTT. The concentration of actin was determined by OD280 and Tetramethylrhodamine-5-Iodoacetamide Dihydroiodide (5-TMRIA) (Fisher Scientific) was added at a 1:1 molar ratio and incubated at 4°C overnight. The following day, 1 mM DTT was added to the mixture, clarified 400,000 x g for 30 minutes and applied to a Superdex 200 Increase 10/300 GL (GE Healthcare) gel filtration column equilibrated in G-buffer containing 0.2 M ammonium acetate. Fractions corresponding to monomeric actin were pooled, concentrated in an Amicon Ultra – 15 (Millipore), snap frozen in liquid nitrogen and stored at –80°C. For generating rhodamine labeled actin filaments, labeled and unlabeled TgAct1 was clarified 400,000 x g for 30 minutes and the concentration was determined by OD280. 1 µM labeled actin was mixed with 20 µM unlabeled actin (5% labeled) and induced for polymerization by adding 2x polymerization buffer without actin chromobody. The mixture was added to a NEM myosin-bound and blocked flow chamber and allowed to polymerize for 10 minutes at 37°C. The flow chamber was then flushed with polymerization buffer containing 50 nM actin chromobody and imaged using epifluorescence microscopy. The resulting actin filaments which were observed using the actin chromobody, produced sparse punctate rhodamine labeling indicating poor rhodamine-TgAct1 incorporation. The average rhodamine-TgAct1 incorporation was determined by counting rhodamine-labeled monomers and dividing by the number of monomers in the filament which was determined using 2.74 nm length increase per subunit (24) (**Ext. Data Fig. 1**).

### Negative Stain Electron Microscopy

Frozen TgAct1 aliquots were thawed at room temperature and then placed on ice. Samples were spun at 95k for 30 min at 4°C and the supernatant was desalted using Zeba spin desalting column (7K MWCO, Thermo Scientific) equilibrated with G-buffer supplemented with 0.2 mM DTT and 0.2 mM Na-ATP. Polymerization was induced by diluting 10x polymerization buffer to 1x in 60 µM (+/-10) TgAct1 with additives depending on the condition. For TgAct1 + jasplakinolide: jasplakinolide was added to a final concentration of 33 µM; for TgAct1 + ATP: 10 mM Mg-ATP was used in the 10x polymerization buffer for a final concentration of 1 mM with TgAct1; for TgAct1 + AMPPNP, Li/Na-AMPPNP was substituted for Mg-ATP in the 10x polymerization buffer for a final concentration of 0.1 mM with TgAct1. Immediately after diluting polymerization buffer into TgAct1, the samples were placed at 37°C and incubated for 1 hour (TgAct1 + Jas), 45 minutes (TgAct1 with AMPPNP), or samples were taken at time points (TgAct1 with 1 mM ATP). Samples were diluted 10-fold in 1x polymerization buffer or directly applied to glow-discharged continuous carbon film grids. Grids were then washed in ddH_2_O three times unless specified otherwise and negatively stained using 2% uranyl formate. Samples were imaged using a FEI Morgagni operating at 100 kV and a Gatan Orius CCD camera with the software package Digital Micrograph (v2.10.1282.0), or using a FEI Tecnai G2 Spirit operating at 120 kV and a Gatan Ultrascan 4000 CCD camera with the software package Leginon (v3.4).

### Cryo-Electron Microscopy & Data Processing

TgAct1 filaments were assembled as described above. TgAct1 + jasplakinolide was assembled and stored at 4°C overnight before grid application. TgAct1 was used immediately after a 20 min incubation at 37°C. Protein solutions were diluted to the concentrations listed in Data Table 1 using 1x polymerization buffer and immediately applied to glow-discharged, C-flat 2/2 holey-carbon EM grids (Protochips Inc.), blotted from the front (TgAct1 + jasplakinolide) or back (TgAct1) of the grid, and plunge-frozen in liquid ethane using a manual plunging apparatus at room temperature in a dehumidified room. Data collection was performed using an FEI Glacios (equipped with a Gatan K3 Summit direct electron detector operating in CDS mode) and an FEI Titan Krios transmission electron microscope operating at 300 kV (equipped with a Gatan image filter (GIF) and post-GIF Gatan K3 Summit direct electron detector) both using the software package Leginon (67) (v3.5).

Movies were aligned, corrected for beam-induced motion and dose-weighted using the Relion (68) 4.0 implementation of MotionCor2 (69) (v1.3.1); CTF parameters were initially estimated using CTFFind4 (v4.1.10) (70). Motion corrected micrographs were imported into cryoSPARC (v4) CTF parameters were re-estimated using Patch CTF Estimation. Particles were picked using Filament Tracer (71). Two rounds of 2D classification were used to remove noise, carbon edges, and poorly aligning particles, and the remaining particles were used in *ab initio* reconstruction and Helix Refinement. Initial helical parameters of –167° twist and 27 Å rise were used (72) within a search space of 164-170° and 24-30 Å; helical parameters were refined in all Helix Refinements. Additional rounds of Helix Refinement with dynamic masks and Local Refinement with a static mask surrounding three protomers of the filament were completed, interspersed with Global and Local CTF Refinement and map sharpening with resolution estimation (FSC cutoff of 0.143). For the TgAct1 dataset, the particles were signal subtracted to remove filament signal outside of the three central protomers prior to the final Local Refinement. The final refined, unsharpened maps were exported to Phenix (v1.20) where density modification and additional resolution estimation was performed (FSC cutoff of 0.5) (73). Local resolution estimation was performed using cryoSPARC’s Local Resolution Estimation and Local Filtering.

### Model Building

PDB ID 6TU4 was used as an initial model for TgAct1. The model was fitted with the correct residues and iteratively refined against both the filament and 3x protomer maps using a combination of ISOLDE (74) (v1.5) in ChimeraX (75) (v1.5), Coot (76) (v0.9.8.1), and phenix.real_space_refine and phenix.validation_cryoem in Phenix (v1.20) (77, 78). The TgAct1 model was used as the initial model for the TgAct1 + jasplakinolide dataset.

### Image Generation and Calculations for Micrographs and Protein Models

Negative stained micrographs were contrast adjusted, rotated, and cropped in Fiji (v2.1.0/1.53c). Cryo electron micrographs were motion corrected in Relion’s implementation of MotionCorr2 and Gaussian filtered and contrast adjusted in Fiji. We used ChimeraX for: protein volume and model visualization, volume alignment (Fit to Model tool or the “fit” command), model alignment (Matchmaker tool or the “align” command), buried surface area calculations (“measure buriedarea” command), hydrophobicity calculations (“hydrophobic coloring” tool), and hydrogen bonds identification (“hbond” tool). All electron micrograph image, volume, and protein model figures were assembled in Adobe Illustrator CC6 (v26.0.1).

## Acknowledgements

We thank Kathleen Trybus and Patricia Fagnant (University of Vermont) for providing DNA subcloning constructs and helpful guidance. We thank our colleagues at the University of Connecticut for sharing equipment; Kenneth Campellone and Jim Cole for the use of their AKTA, and Victoria Robinson and Giancarlo Montovano their assistance with ion exchange chromatography. We thank the members of the De La Cruz lab (Yale University), Heaslip lab (University of Connecticut) and John Murray (Arizona State University) for thoughtful discussions. We thank Joel Quispe, Sasha Dickinson, and the Arnold and Mabel Beckman Cryo-EM Center at the University of Washington for electron microscope guidance and use. We also thank members of the Kollman group for feedback provided during cryo-EM data collection and processing. Thanks to Alexander Paredez for initiating this fruitful collaboration. This work was supported by the National Institutes of General Medical Science and Allergy and Infectious Disease of the US National Institutes of Health (grant nos. R35GM136656 to E.M.D.L.C., R35GM149542 and S10OD023476 to J.M.K., 1F32AI145111 to K.L.H., and R35GM138316 to A.T.H.).

## Author Contributions Statement

T.E.S. purified TgAct1 and actin chromobody, designed and performed actin growth and biochemical assays. Kinetic assays were designed by E.M.D.L.C and performed by T.E.S. K.L.H. designed and performed negative stain and cryo electron microscopy experiments. T.E.S., K.L.H., J.M.K., and A.T.H. performed data analysis and interpretation and wrote the manuscript. All authors edited and revised the manuscript.

## Competing Interests Statement

The authors declare no competing interests.

## Supplementary Figures

**Ext. Data Figure 1:**
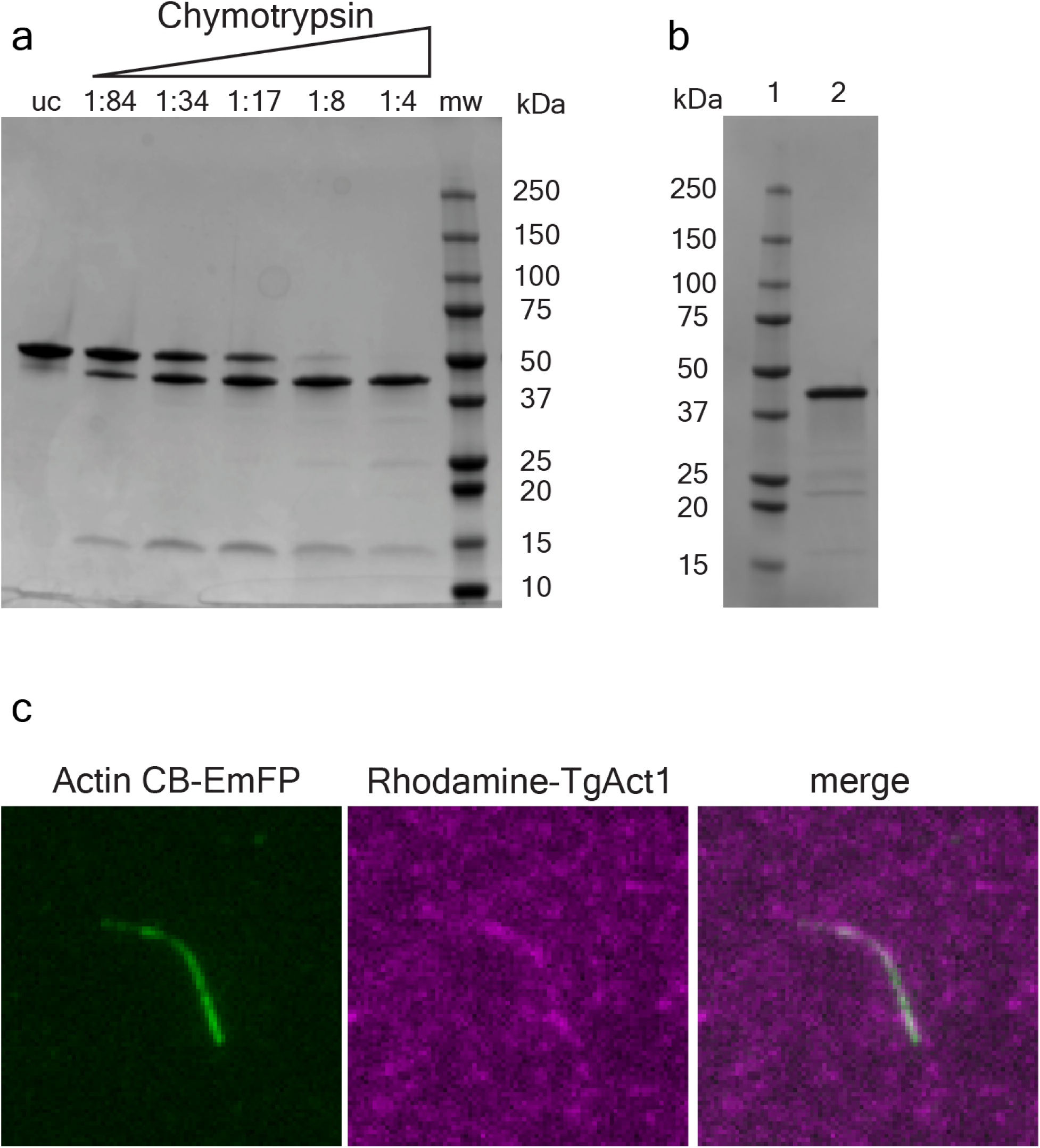
Protein preparations and growth of TgAct1 in the presence of labeled Rhodamine actin. **a**, Coomassie stained SDS-PAGE showing TgAct1–β-thymosin–HIS incubated with different molar ratios of chymotrypsin:actin for 15 min at 27°C, to determine optimal conditions for removal of β-thymosin–HIS tag. uc: uncut; mw: protein molecular weight marker. **b,** Coomassie stained SDS-PAGE gel showing (lane 1) protein molecular weight marker and (lane 2) 2 µg purified actin chromobody-EmeraldFP. **c,** Unlabeled TgAct1 was mixed with Rhodamine-labeled TgAct1 at a ratio of 20:1 (5%) and induced for polymerization for 15 minutes. The resulting filaments were adhered to a flow chamber, washed with an imaging buffer containing actin chromobody-EmeraldFP and imaged. The average percent incorporation (0.5 ± 0.07%, n = 4 filaments) was determined by visualizing the number of Rhodamine positive monomers per filament length.

**Ext. Data Figure 2:**
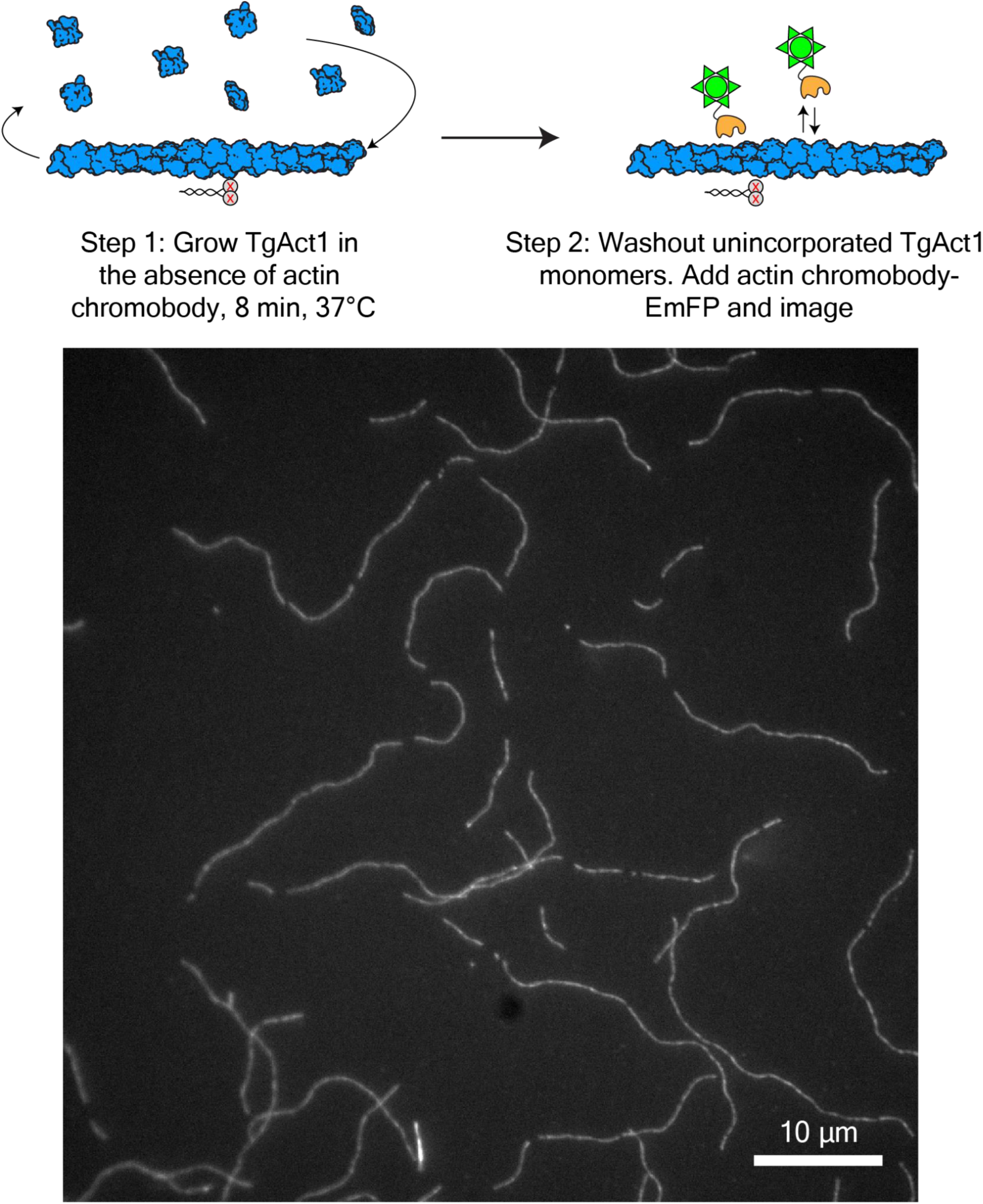
Imaging TgAct1 polymerized in the absence of chromobody. TgAct1 was induced for polymerization in a flow chamber in the absence of actin chromobody for 10 min at 37°C. Then, unincorporated TgAct1 monomers were washed out and replaced with a solution containing 50 nM actin chromobody-EmeraldFP and imaged before filaments depolymerized. Polymerization conditions: 25 mM imidazole, pH 7.4, 50 mM KCl, 2.5 mM MgCl_2_, 1 mM EGTA, 2.5 mM MgATP,10 mM DTT, 0.25% methylcellulose, 2.5 mg/mL BSA, 0.5% Pluronic F-127, oxygen scavenging system (0.13 mg/mL glucose oxidase, 50 μg/mL catalase, and 3 mg/mL glucose), 37°C.

**Ext. Data Figure 3:**
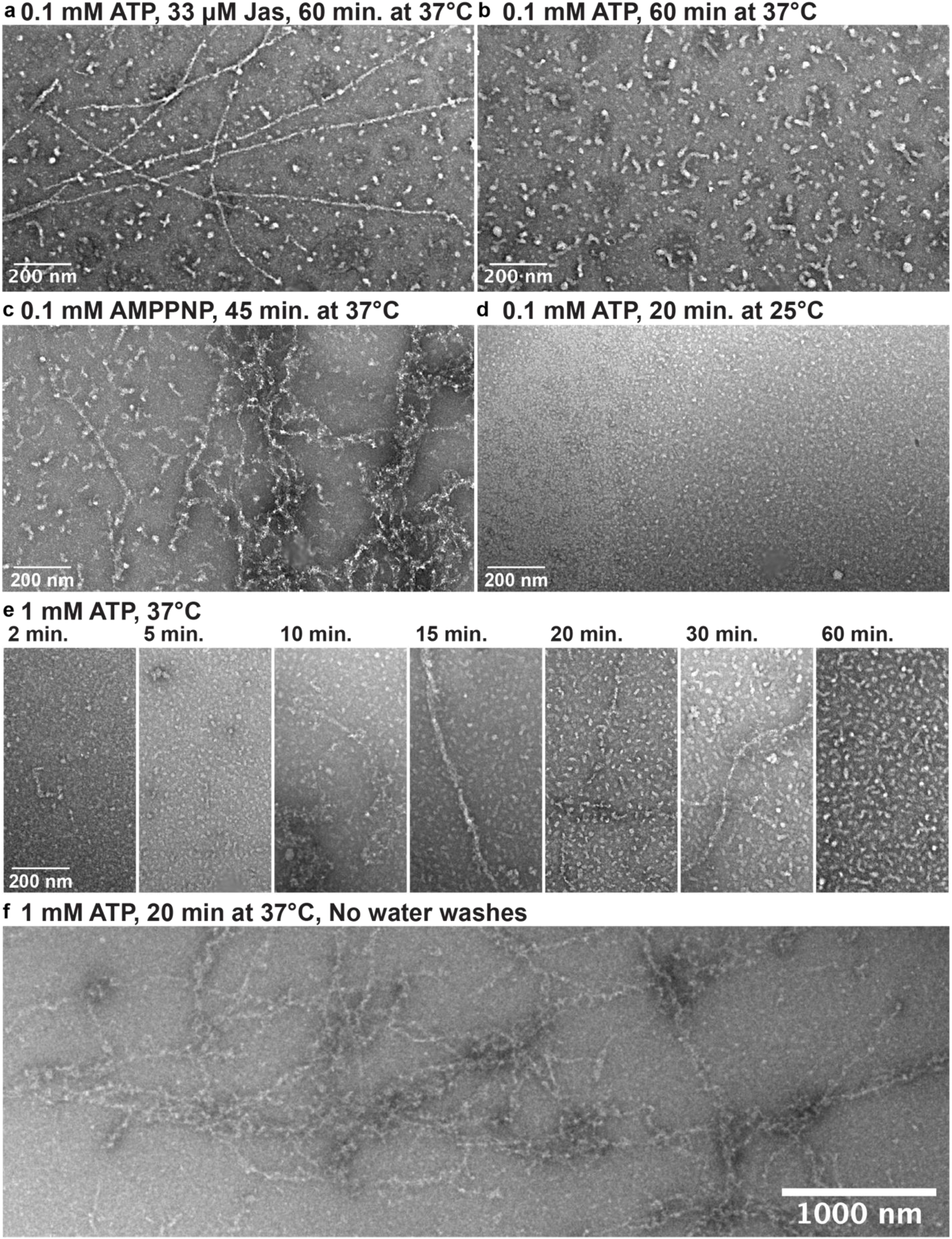
Negative stain electron microscopy of TgAct1 filaments. **a**, After 1 hour at 37°C in polymerization buffer + jasplakinolide, filaments are observed. **b,** Under the same conditions but without jasplakinolide, no filaments are observed. **c,** Substitution of AMPPNP for Mg-ATP produces observable filaments. **d,** No filaments were observed by shortening the incubation time and a decrease in the temperature. **e,** Filaments were observed in conditions where the concentration of Mg-ATP was increased and shorter time points were checked (10-30 min). **f,** Increased numbers and lengths of filaments were identified on grids where the water washes in between protein application and addition of uranyl formate were skipped.

**Ext. Data Figure 4:**
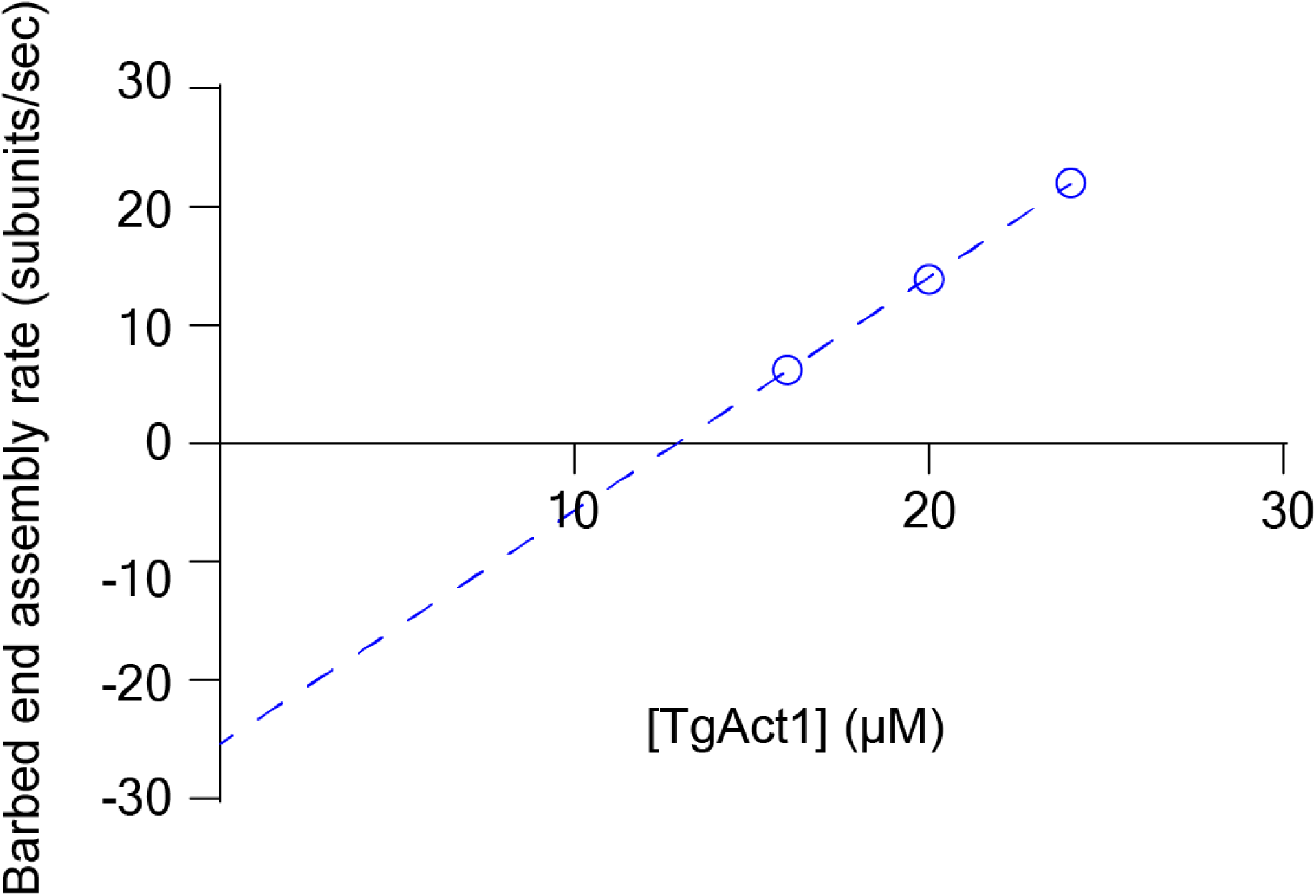
Plot of the rate of barbed-end growth in subunits/sec, per actin concentration for TgAct1 in the presence of 0.1 mM AMPPNP. The critical concentration (*C_c_*) (x-intercept of the fitted line) is 12.9 µM. The dissociation constant (k_-_) (y-intercept) is 25.4 sec^-1^ and the association constant (k_+_) (slope) is 2.0 sec^-1^ µM^-1^.

**Ext. Data Figure 5:**
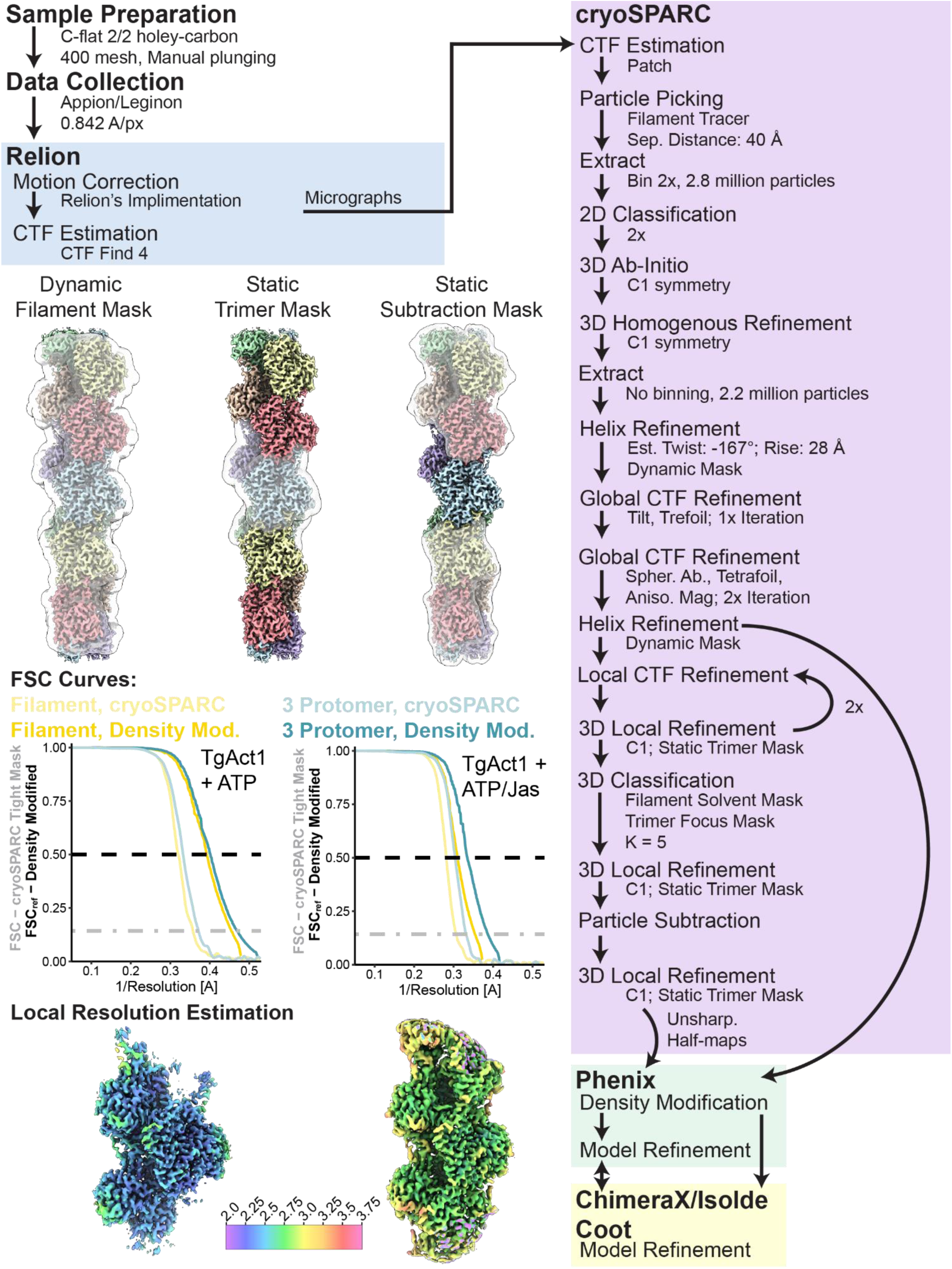
Data processing scheme and statistics for cryo electron microscopy datasets. The flowchart shows the data processing scheme for TgAct1 + 1 mM ATP; a similar scheme without particle subtraction was used for TgAct1 + jasplakinolide. The masks shown were used as described in the flowchart. Fourier shell correlation curves for the helical reconstructions are shown in yellow and curves for the locally-refined, three-protomer volume are shown in blue, shaded based on the software in which they were calculated (cryoSPARC, light colors; Phenix Density Modification, dark colors). Local resolutions estimated in cryoSPARC were mapped onto volumes locally filtered in cryoSPARC.

**Ext. Data Figure 6:**
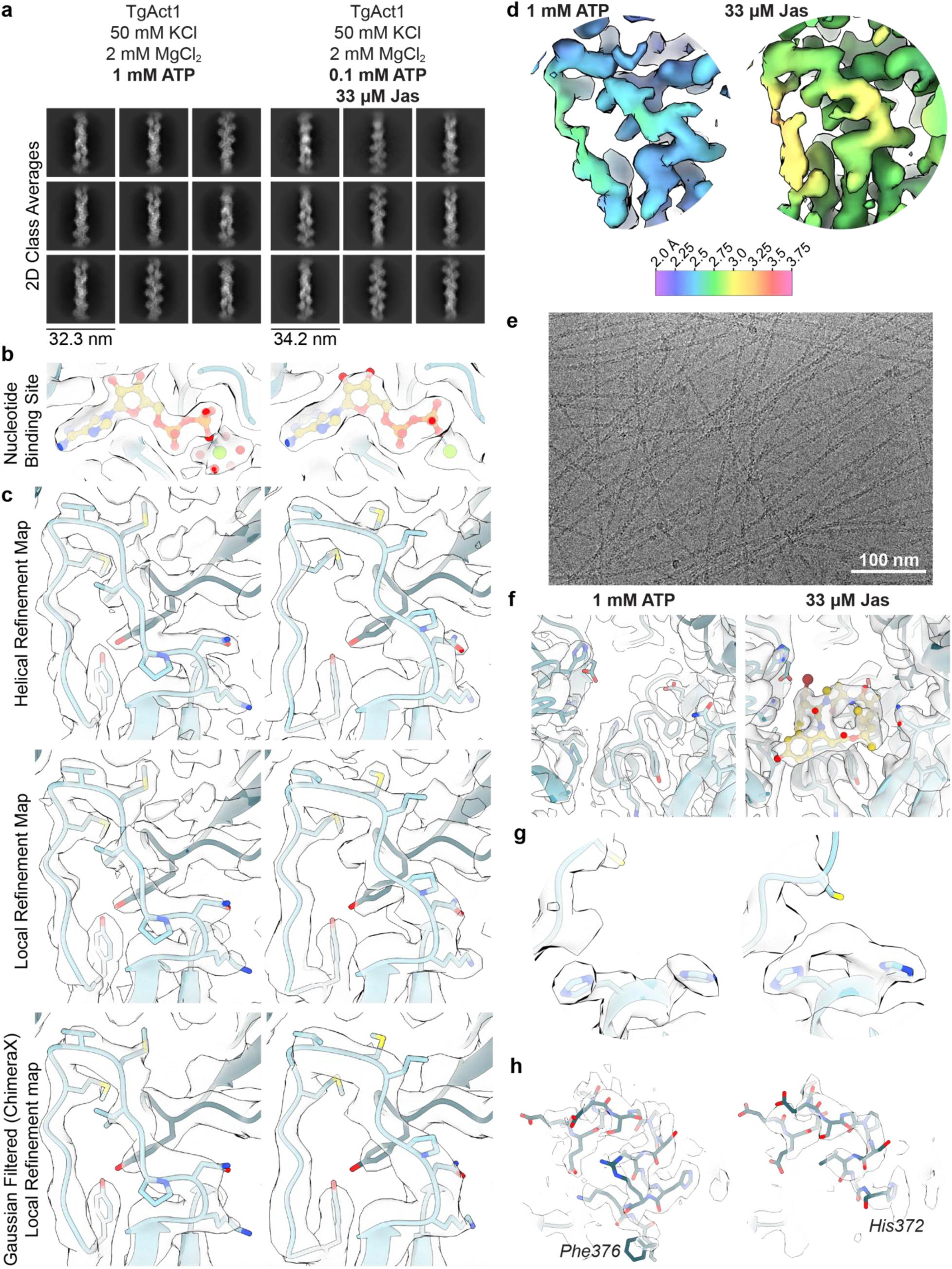
Exploration of TgAct1 filament data, maps, and models. **a**, Representative two-dimensional class averages from the data sets. **b,** Map and model from each dataset showing the nucleotide binding site, with the protein in blue and the nucleotide (ADP in both structures) in yellow. **c,** Volumes from helical refinement (top) and local refinement (middle) maps. The bottom panels show the local refinement map after Gaussian filtering in ChimeraX. All models are the final model for the dataset indicated. **d,** Local resolution maps of the D-loop for each filament (unstabilized TgAct, left and TgAct1 + jasplakinolide, right). **e,** Representative micrograph from the TgAct1 + jasplakinolide dataset. **f,** Map and model from each dataset showing the jasplakinolide binding site (unstabilized TgAct, left and TgAct1 + jasplakinolide, right). **g,** View of Cys53 from each dataset showing volume that suggests the cysteine has been oxidized. **h,** Volumes and models of the C-termini from each data set.

**Ext. Data Figure 7:**
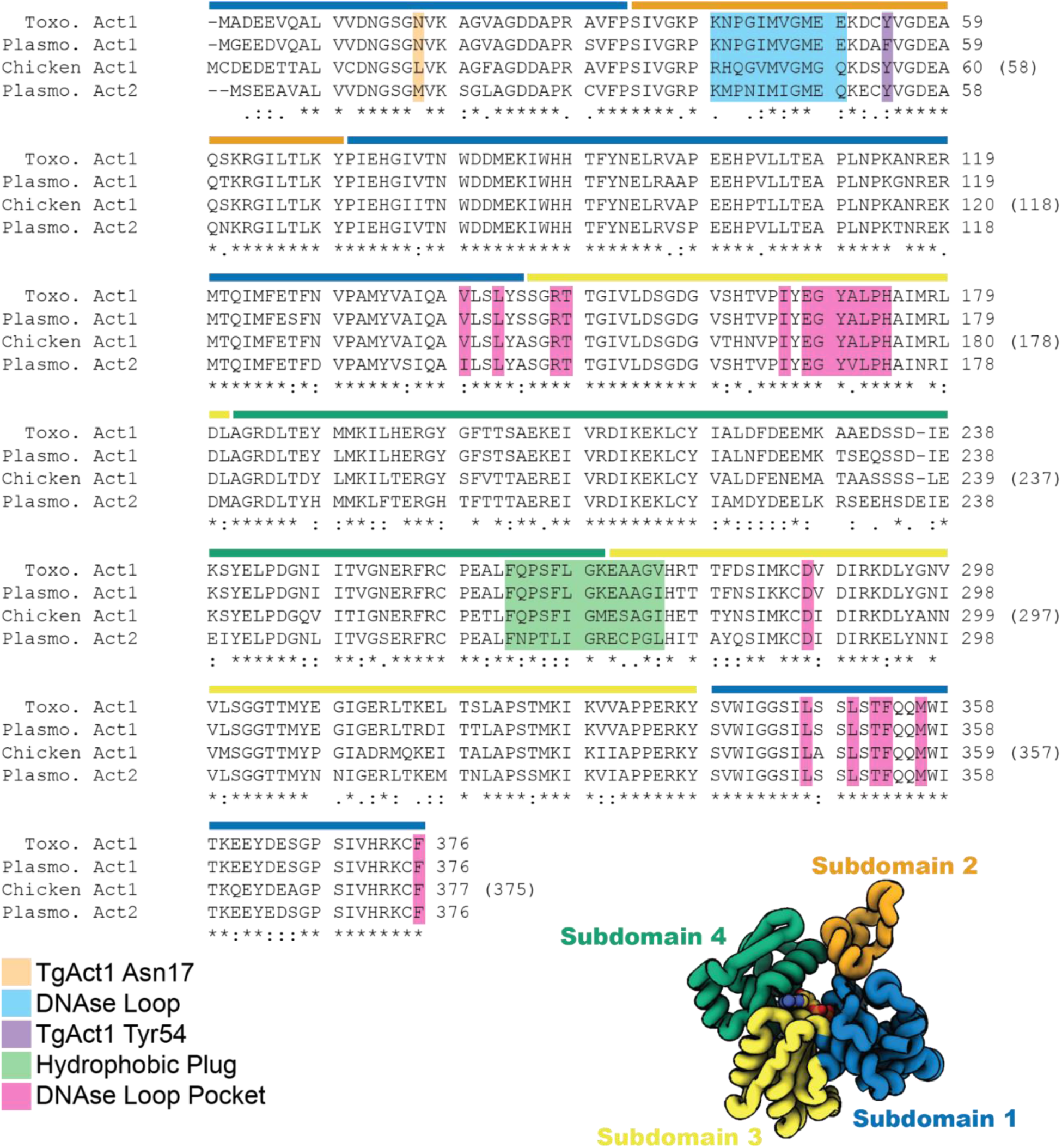
Sequence alignment. Sequence alignment of TgAct1, skeletal actin, *P. falciparum* Act1, and *P. falciparum* Act2. Sequences retrieved from Uniprot (Uniprot IDs: XYZ) and aligned with the EMBL-EBI portal for MAFFT (v7, https://www.ebi.ac.uk/Tools/msa/mafft/).

**Ext. Data Figure 8:**
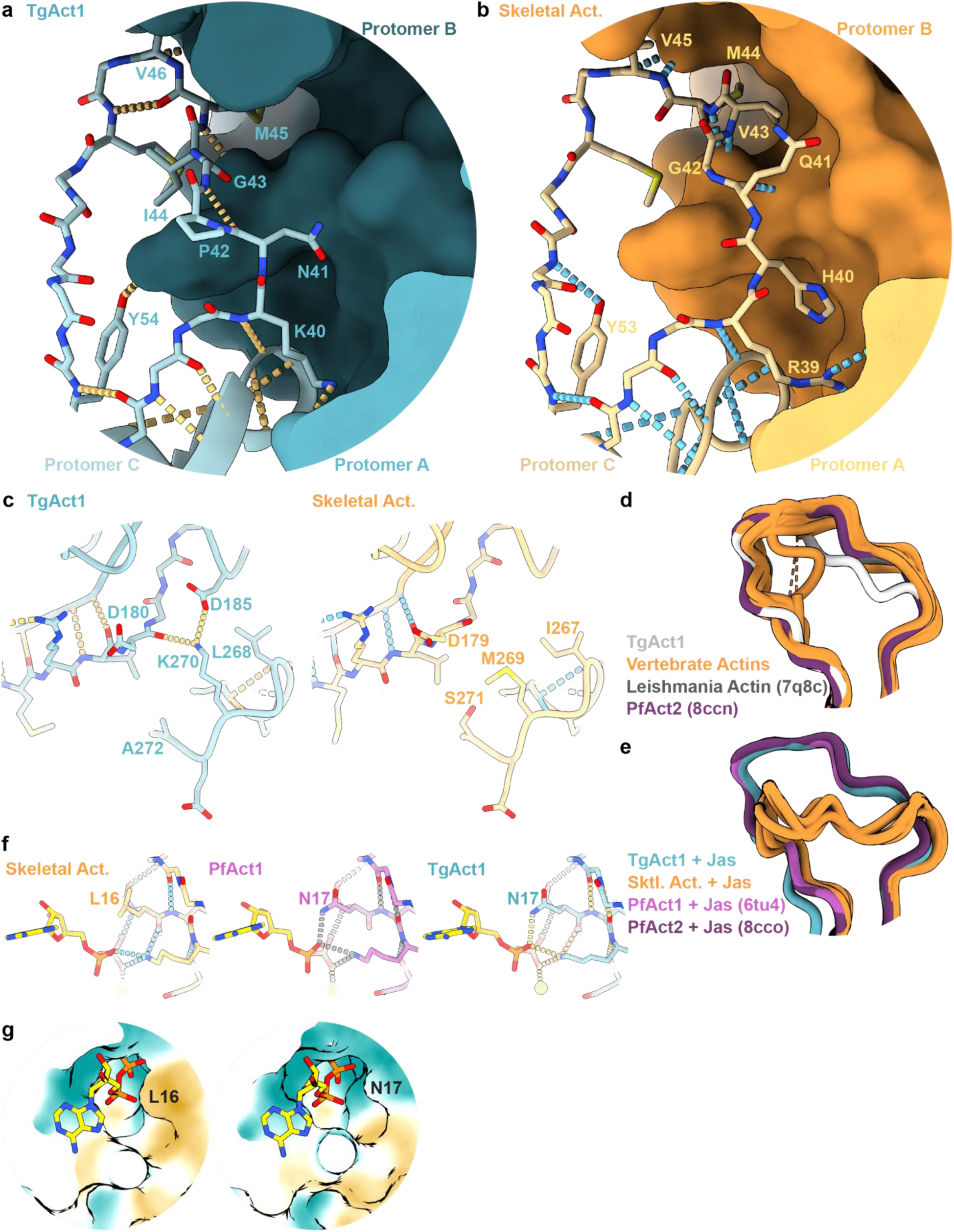
Actin filament comparisons. **a/b**, Hydrogen bonding network of D-loop from unstabilized TgAct1 actin filaments (protein in blue, hydrogen bonds in orange) and skeletal actin filaments (protein in orange, hydrogen bonds in blue). Side chains of amino acids 49-53 (skeletal actin 48-52) have been omitted for clarity. **c,** Hydrogen bonding network of the hydrophobic plug region of unstabilized TgAct1 actin filaments (protein in blue, hydrogen bonds in orange) and skeletal actin filaments (protein in orange, hydrogen bonds in blue). **d,** The D-loops from unstabilized TgAct1 (gray), several vertebrate actins (orange, PDB IDs 7bt7, 7r8v, 8a2r, 8a2t, 8d13, 8dmx, 8dmy, 8dnf, 8dnh), Leishmania actin (dark gray, PDB ID 7q8c), and *P. falciparum* Act2 (purple, PDB ID 8ccn). **e,** The D-loops from TgAct1 + jasplakinolide (blue), skeletal actin + jasplakinolide (orange, PDB IDs 5ooc, 5ood, 6t24, 7pm3), *P. falciparum* Act1 + jasplakinolide (magenta, PDB ID 6tu4), and *P. falciparum* Act2 + jasplakinolide (purple, PDB ID 8cco). **f,** View of Leu16 or Asn17 from the nucleotide binding pocket of skeletal actin filaments (orange, hydrogen bonds in blue), *P. falciparum* Act1 filaments (magenta, hydrogen bonds in grey), and unstabilized TgAct1 (blue, hydrogen bonds in orange). **g,** Surface representation of the nucleotide binding pocket of skeletal G-actin (PDB ID 1eqy) and *P. falciparum* Act1 G-actin (PDB ID 6i4e) with the nucleotide shown as sticks. Surface colors represent hydrophobicity, with oranges being the most hydrophobic and teals being the most hydrophilic.

## Supplementary information

**Supplementary Video 1:** Direct real-time visualization of TgAct1 polymerization in the presence of 16 µM TgAct1 and 25 nM actin chromobody-EmeraldFP. Conditions: 25 mM imidazole, pH 7.4, 50 mM KCl, 2.5 mM MgCl_2_, 1 mM EGTA, 2.5 mM MgATP,10 mM DTT, 0.25% methylcellulose, 2.5 mg/mL BSA, 0.5% Pluronic F-127, oxygen scavenging system (0.13 mg/mL glucose oxidase, 50 μg/mL catalase, and 3 mg/mL glucose), 37°C. 100x playback, Image width 66 µm.

**Supplementary Video 2:** Example of treadmilling TgAct1 filaments in the presence of 16 µM TgAct1 and 25 nM actin chromobody-EmeraldFP. 100x playback, Image width 21 µm.

**Supplementary Video 3**: Ribbon morph between an unstabilized TgAct1 filament and skeletal actin filament. Movie morphs from unstabilized TgAct1 filament to the skeletal muscle actin filament (8d13) and returns to the unstabilized TgAct1 filament.

**Supplementary Video 4**: Ribbon morph between an unstabilized TgAct1 filament and jasplakinolide-bound TgAct1 filaments. Movie morphs from unstabilized TgAct1 filament to the jasplakinolide-bound TgAct1 and returns to the unstabilized TgAct1 filament. Jasplakinolide shown in yellow for reference.

## Data availability

The cryo-EM maps generated for this manuscript are available from the EMDB (https://www.ebi.ac.uk/emdb/) at the accession codes listed in Table 1 of the manuscript (EMDB IDs: XYZ1, XYZ2) The protein models generated for this manuscript are available from the RCSB PDB (https://www.rcsb.org/) at the accession codes listed in Table 1 of the manuscript (PDB IDs: XYZ1, XYZ2). Protein sequences were retrieved from Uniprot (https://www.uniprot.org/) at the codes: P53476, Q8I4X0, P68139, and Q8ILW9. Source data are provided with this paper.

## Notes

### Competing Interest Statement

The authors have declared no competing interest.

## References

1. Rorman E, Zamir CS, Rilkis I, Ben-David H. Congenital toxoplasmosis--prenatal aspects of Toxoplasma gondii infection. Reprod Toxicol. 2006;21(4):458–72. doi: 10.1016/j.reprotox.2005.10.006. PubMed PMID: 16311017.

2. Wang ZD, Wang SC, Liu HH, Ma HY, Li ZY, Wei F, Zhu XQ, Liu Q. Prevalence and burden of Toxoplasma gondii infection in HIV-infected people: a systematic review and meta-analysis. Lancet HIV. 2017;4(4):e177–e88. doi: 10.1016/S2352-3018(17)30005-X. PubMed PMID: 28159548.

3. Torgerson PR, Mastroiacovo P. The global burden of congenital toxoplasmosis: a systematic review. Bull World Health Organ. 2013;91(7):501–8. doi: 10.2471/BLT.12.111732. PubMed PMID: 23825877; PMCID: PMC3699792.

4. Dobrowolski JM, Niesman IR, Sibley LD. Actin in the parasite Toxoplasma gondii is encoded by a single copy gene, ACT1 and exists primarily in a globular form. Cell Motil Cytoskeleton. 1997;37(3):253–62. doi: 10.1002/(SICI)1097-0169(1997)37:3<253::AID-CM7>3.0.CO;2-7. PubMed PMID: 9227855.

5. Dobrowolski JM, Sibley LD. Toxoplasma invasion of mammalian cells is powered by the actin cytoskeleton of the parasite. Cell. 1996;84(6):933–9. doi: 10.1016/s0092-8674(00)81071-5. PubMed PMID: 8601316.

6. Poupel O, Tardieux I. Toxoplasma gondii motility and host cell invasiveness are drastically impaired by jasplakinolide, a cyclic peptide stabilizing F-actin. Microbes Infect. 1999;1(9):653–62. doi: 10.1016/s1286-4579(99)80066-5. PubMed PMID: 10611742.

7. Mehta S, Sibley LD. Actin depolymerizing factor controls actin turnover and gliding motility in Toxoplasma gondii. Mol Biol Cell. 2011;22(8):1290–9. doi: 10.1091/mbc.E10-12-0939. PubMed PMID: 21346192; PMCID: PMC3078074.

8. Whitelaw JA, Latorre-Barragan F, Gras S, Pall GS, Leung JM, Heaslip A, Egarter S, Andenmatten N, Nelson SR, Warshaw DM, Ward GE, Meissner M. Surface attachment, promoted by the actomyosin system of Toxoplasma gondii is important for efficient gliding motility and invasion. BMC Biol. 2017;15(1):1. doi: 10.1186/s12915-016-0343-5. PubMed PMID: 28100223; PMCID: PMC5242020.

9. Jacot D, Daher W, Soldati-Favre D. Toxoplasma gondii myosin F, an essential motor for centrosomes positioning and apicoplast inheritance. EMBO J. 2013;32(12):1702–16. doi: 10.1038/emboj.2013.113. PubMed PMID: 23695356; PMCID: PMC3680736.

10. Devarakonda PM, Sarmiento V, Heaslip AT. F-actin and Myosin F control apicoplast elongation dynamics which drive apicoplast-centrosome association in Toxoplasma gondii. bioRxiv. 2023. doi: 10.1101/2023.01.01.521342. PubMed PMID: 36711828; PMCID: PMC9881852.

11. Heaslip AT, Nelson SR, Warshaw DM. Dense granule trafficking in Toxoplasma gondii requires a unique class 27 myosin and actin filaments. Mol Biol Cell. 2016;27(13):2080–9. doi: 10.1091/mbc.E15-12-0824. PubMed PMID: 27146112; PMCID: PMC4927281.

12. Periz J, Del Rosario M, McStea A, Gras S, Loney C, Wang L, Martin-Fernandez ML, Meissner M. A highly dynamic F-actin network regulates transport and recycling of micronemes in Toxoplasma gondii vacuoles. Nat Commun. 2019;10(1):4183. doi: 10.1038/s41467-019-12136-2. PubMed PMID: 31519913; PMCID: PMC6744512.

13. Carmeille R, Schiano Lomoriello P, Devarakonda PM, Kellermeier JA, Heaslip AT. Actin and an unconventional myosin motor, TgMyoF, control the organization and dynamics of the endomembrane network in Toxoplasma gondii. PLoS Pathog. 2021;17(2):e1008787. doi: 10.1371/journal.ppat.1008787. PubMed PMID: 33529198; PMCID: PMC7880465.

14. Dos Santos Pacheco N, Brusini L, Haase R, Tosetti N, Maco B, Brochet M, Vadas O, Soldati-Favre D. Conoid extrusion regulates glideosome assembly to control motility and invasion in Apicomplexa. Nat Microbiol. 2022;7(11):1777–90. doi: 10.1038/s41564-022-01212-x. PubMed PMID: 36109645.

15. Das S, Stortz JF, Meissner M, Periz J. The multiple functions of actin in apicomplexan parasites. Cell Microbiol. 2021;23(11):e13345. doi: 10.1111/cmi.13345. PubMed PMID: 33885206.

16. Periz J, Whitelaw J, Harding C, Gras S, Del Rosario Minina MI, Latorre-Barragan F, Lemgruber L, Reimer MA, Insall R, Heaslip A, Meissner M. Toxoplasma gondii F-actin forms an extensive filamentous network required for material exchange and parasite maturation. Elife. 2017;6. doi: 10.7554/eLife.24119. PubMed PMID: 28322189; PMCID: PMC5375643.

17. Stortz JF, Del Rosario M, Singer M, Wilkes JM, Meissner M, Das S. Formin-2 drives polymerisation of actin filaments enabling segregation of apicoplasts and cytokinesis in Plasmodium falciparum. Elife. 2019;8. doi: 10.7554/eLife.49030. PubMed PMID: 31322501; PMCID: PMC6688858.

18. Tosetti N, Dos Santos Pacheco N, Soldati-Favre D, Jacot D. Three F-actin assembly centers regulate organelle inheritance, cell-cell communication and motility in Toxoplasma gondii. Elife. 2019;8. doi: 10.7554/eLife.42669. PubMed PMID: 30753127; PMCID: PMC6372287.

19. Panza P, Maier J, Schmees C, Rothbauer U, Sollner C. Live imaging of endogenous protein dynamics in zebrafish using chromobodies. Development. 2015;142(10):1879–84. doi: 10.1242/dev.118943. PubMed PMID: 25968318; PMCID: PMC4440926.

20. Melak M, Plessner M, Grosse R. Actin visualization at a glance. J Cell Sci. 2017;130(3):525–30. doi: 10.1242/jcs.189068. PubMed PMID: 28082420.

21. Lu H, Fagnant PM, Trybus KM. Unusual dynamics of the divergent malaria parasite PfAct1 actin filament. Proc Natl Acad Sci U S A. 2019;116(41):20418–27. doi: 10.1073/pnas.1906600116. PubMed PMID: 31548388; PMCID: PMC6789906.

22. Skillman KM, Ma CI, Fremont DH, Diraviyam K, Cooper JA, Sept D, Sibley LD. The unusual dynamics of parasite actin result from isodesmic polymerization. Nat Commun. 2013;4:2285. doi: 10.1038/ncomms3285. PubMed PMID: 23921463; PMCID: PMC3765016.

23. Kumpula EP, Pires I, Lasiwa D, Piirainen H, Bergmann U, Vahokoski J, Kursula I. Apicomplexan actin polymerization depends on nucleation. Sci Rep. 2017;7(1):12137. doi: 10.1038/s41598-017-11330-w. PubMed PMID: 28939886; PMCID: PMC5610305.

24. Vahokoski J, Bhargav SP, Desfosses A, Andreadaki M, Kumpula EP, Martinez SM, Ignatev A, Lepper S, Frischknecht F, Siden-Kiamos I, Sachse C, Kursula I. Structural differences explain diverse functions of Plasmodium actins. PLoS Pathog. 2014;10(4):e1004091. doi: 10.1371/journal.ppat.1004091. PubMed PMID: 24743229; PMCID: PMC3990709.

25. Bookwalter CS, Tay CL, McCrorie R, Previs MJ, Lu H, Krementsova EB, Fagnant PM, Baum J, Trybus KM. Reconstitution of the core of the malaria parasite glideosome with recombinant Plasmodium class XIV myosin A and Plasmodium actin. J Biol Chem. 2017;292(47):19290–303. doi: 10.1074/jbc.M117.813972. PubMed PMID: 28978649; PMCID: PMC5702669.

26. Noguchi TQ, Kanzaki N, Ueno H, Hirose K, Uyeda TQ. A novel system for expressing toxic actin mutants in Dictyostelium and purification and characterization of a dominant lethal yeast actin mutant. J Biol Chem. 2007;282(38):27721–7. doi: 10.1074/jbc.M703165200. PubMed PMID: 17656358.

27. Fujiwara I, Zweifel ME, Courtemanche N, Pollard TD. Latrunculin A Accelerates Actin Filament Depolymerization in Addition to Sequestering Actin Monomers. Curr Biol. 2018;28(19):3183–92 e2. doi: 10.1016/j.cub.2018.07.082. PubMed PMID: 30270183; PMCID: PMC6179359.

28. Baum J, Papenfuss AT, Baum B, Speed TP, Cowman AF. Regulation of apicomplexan actin-based motility. Nat Rev Microbiol. 2006;4(8):621–8. doi: 10.1038/nrmicro1465. PubMed PMID: 16845432.

29. Kuhn JR, Pollard TD. Real-time measurements of actin filament polymerization by total internal reflection fluorescence microscopy. Biophys J. 2005;88(2):1387–402. doi: 10.1529/biophysj.104.047399. PubMed PMID: 15556992; PMCID: PMC1305141.

30. Pollard TD. Rate constants for the reactions of ATP– and ADP-actin with the ends of actin filaments. J Cell Biol. 1986;103(6 Pt 2):2747–54. doi: 10.1083/jcb.103.6.2747. PubMed PMID: 3793756; PMCID: PMC2114620.

31. Melki R, Fievez S, Carlier MF. Continuous monitoring of Pi release following nucleotide hydrolysis in actin or tubulin assembly using 2-amino-6-mercapto-7-methylpurine ribonucleoside and purine-nucleoside phosphorylase as an enzyme-linked assay. Biochemistry. 1996;35(37):12038–45. doi: 10.1021/bi961325o. PubMed PMID: 8810908.

32. Blanchoin L, Pollard TD. Mechanism of interaction of Acanthamoeba actophorin (ADF/Cofilin) with actin filaments. J Biol Chem. 1999;274(22):15538–46. doi: 10.1074/jbc.274.22.15538. PubMed PMID: 10336448.

33. Korn ED, Carlier MF, Pantaloni D. Actin polymerization and ATP hydrolysis. Science. 1987;238(4827):638-44. doi: 10.1126/science.3672117. PubMed PMID: 3672117.

34. Carlier MF, Pantaloni D, Evans JA, Lambooy PK, Korn ED, Webb MR. The hydrolysis of ATP that accompanies actin polymerization is essentially irreversible. FEBS Lett. 1988;235(1-2):211–4. doi: 10.1016/0014-5793(88)81264-x. PubMed PMID: 2969829.

35. Carlier MF. Role of nucleotide hydrolysis in the dynamics of actin filaments and microtubules. Int Rev Cytol. 1989;115:139–70. doi: 10.1016/s0074-7696(08)60629-4. PubMed PMID: 2663760.

36. Pollard TD. Actin. Curr Opin Cell Biol. 1990;2(1):33–40. doi: 10.1016/s0955-0674(05)80028-6. PubMed PMID: 2183841.

37. Kinosian HJ, Selden LA, Estes JE, Gershman LC. Nucleotide binding to actin. Cation dependence of nucleotide dissociation and exchange rates. J Biol Chem. 1993;268(12):8683–91. PubMed PMID: 8473312.

38. Pollard TD, Goldberg I, Schwarz WH. Nucleotide exchange, structure, and mechanical properties of filaments assembled from ATP-actin and ADP-actin. J Biol Chem. 1992;267(28):20339–45. PubMed PMID: 1400353.

39. Frieden C, Patane K. Mechanism for nucleotide exchange in monomeric actin. Biochemistry. 1988;27(10):3812–20. doi: 10.1021/bi00410a044. PubMed PMID: 3408729.

40. Neidl C, Engel J. Exchange of ADP, ATP and 1: N6-ethenoadenosine 5’-triphosphate at G-actin. Equilibrium and kinetics. Eur J Biochem. 1979;101(1):163–9. doi: 10.1111/j.1432-1033.1979.tb04228.x. PubMed PMID: 510301.

41. De La Cruz EM, Pollard TD. Nucleotide-free actin: stabilization by sucrose and nucleotide binding kinetics. Biochemistry. 1995;34(16):5452–61. doi: 10.1021/bi00016a016. PubMed PMID: 7727403.

42. Mockrin SC, Korn ED. Acanthamoeba profilin interacts with G-actin to increase the rate of exchange of actin-bound adenosine 5’-triphosphate. Biochemistry. 1980;19(23):5359–62. doi: 10.1021/bi00564a033. PubMed PMID: 6893804.

43. Nishida E. Opposite effects of cofilin and profilin from porcine brain on rate of exchange of actin-bound adenosine 5’-triphosphate. Biochemistry. 1985;24(5):1160–4. doi: 10.1021/bi00326a015. PubMed PMID: 4096896.

44. Goldschmidt-Clermont PJ, Machesky LM, Doberstein SK, Pollard TD. Mechanism of the interaction of human platelet profilin with actin. J Cell Biol. 1991;113(5):1081–9. doi: 10.1083/jcb.113.5.1081. PubMed PMID: 1645736; PMCID: PMC2289016.

45. Waechter F, Engel J. The kinetics of the exchange of G-actin-bound 1: N6-ethenoadenosine 5’-triphosphate with ATP as followed by fluorescence. Eur J Biochem. 1975;57(2):453–9. doi: 10.1111/j.1432-1033.1975.tb02320.x. PubMed PMID: 240724.

46. Sahoo N, Beatty W, Heuser J, Sept D, Sibley LD. Unusual kinetic and structural properties control rapid assembly and turnover of actin in the parasite Toxoplasma gondii. Mol Biol Cell. 2006;17(2):895–906. doi: 10.1091/mbc.e05-06-0512. PubMed PMID: 16319175; PMCID: PMC1356598.

47. Reynolds MJ, Hachicho C, Carl AG, Gong R, Alushin GM. Bending forces and nucleotide state jointly regulate F-actin structure. Nature. 2022;611(7935):380–6. doi: 10.1038/s41586-022-05366-w. PubMed PMID: 36289330; PMCID: PMC9646526.

48. Oosterheert W, Klink BU, Belyy A, Pospich S, Raunser S. Structural basis of actin filament assembly and aging. Nature. 2022;611(7935):374–9. doi: 10.1038/s41586-022-05241-8. PubMed PMID: 36289337; PMCID: PMC9646518.

49. Pospich S, Kumpula EP, von der Ecken J, Vahokoski J, Kursula I, Raunser S. Near-atomic structure of jasplakinolide-stabilized malaria parasite F-actin reveals the structural basis of filament instability. Proc Natl Acad Sci U S A. 2017;114(40):10636–41. doi: 10.1073/pnas.1707506114. PubMed PMID: 28923924; PMCID: PMC5635891.

50. Lopez AJ, Andreadaki M, Vahokoski J, Deligianni E, Calder LJ, Camerini S, Freitag A, Bergmann U, Rosenthal PB, Siden-Kiamos I, Kursula I. Structure and function of Plasmodium actin II in the parasite mosquito stages. PLoS Pathog. 2023;19(3):e1011174. doi: 10.1371/journal.ppat.1011174. PubMed PMID: 36877739; PMCID: PMC10019781.

51. Kumpula EP, Lopez AJ, Tajedin L, Han H, Kursula I. Atomic view into Plasmodium actin polymerization, ATP hydrolysis, and fragmentation. PLoS Biol. 2019;17(6):e3000315. doi: 10.1371/journal.pbio.3000315. PubMed PMID: 31199804; PMCID: PMC6599135.

52. McLaughlin PJ, Gooch JT, Mannherz HG, Weeds AG. Structure of gelsolin segment 1-actin complex and the mechanism of filament severing. Nature. 1993;364(6439):685-92. doi: 10.1038/364685a0. PubMed PMID: 8395021.

53. Jegou A, Niedermayer T, Orban J, Didry D, Lipowsky R, Carlier MF, Romet-Lemonne G. Individual actin filaments in a microfluidic flow reveal the mechanism of ATP hydrolysis and give insight into the properties of profilin. PLoS Biol. 2011;9(9):e1001161. doi: 10.1371/journal.pbio.1001161. PubMed PMID: 21980262; PMCID: PMC3181223.

54. Kovar DR, Harris ES, Mahaffy R, Higgs HN, Pollard TD. Control of the assembly of ATP– and ADP-actin by formins and profilin. Cell. 2006;124(2):423–35. doi: 10.1016/j.cell.2005.11.038. PubMed PMID: 16439214.

55. Selden LA, Kinosian HJ, Estes JE, Gershman LC. Impact of profilin on actin-bound nucleotide exchange and actin polymerization dynamics. Biochemistry. 1999;38(9):2769–78. doi: 10.1021/bi981543c. PubMed PMID: 10052948.

56. Bamburg JR. Proteins of the ADF/cofilin family: essential regulators of actin dynamics. Annu Rev Cell Dev Biol. 1999;15:185–230. doi: 10.1146/annurev.cellbio.15.1.185. PubMed PMID: 10611961.

57. Carlier MF, Ressad F, Pantaloni D. Control of actin dynamics in cell motility. Role of ADF/cofilin. J Biol Chem. 1999;274(48):33827–30. doi: 10.1074/jbc.274.48.33827. PubMed PMID: 10567336.

58. Ono S. Regulation of actin filament dynamics by actin depolymerizing factor/cofilin and actin-interacting protein 1: new blades for twisted filaments. Biochemistry. 2003;42(46):13363–70. doi: 10.1021/bi034600x. PubMed PMID: 14621980.

59. Paavilainen VO, Bertling E, Falck S, Lappalainen P. Regulation of cytoskeletal dynamics by actin-monomer-binding proteins. Trends Cell Biol. 2004;14(7):386–94. doi: 10.1016/j.tcb.2004.05.002. PubMed PMID: 15246432.

60. Schuler H, Mueller AK, Matuschewski K. A Plasmodium actin-depolymerizing factor that binds exclusively to actin monomers. Mol Biol Cell. 2005;16(9):4013–23. doi: 10.1091/mbc.e05-02-0086. PubMed PMID: 15975905; PMCID: PMC1196315.

61. Ozyamak E, Kollman J, Agard DA, Komeili A. The bacterial actin MamK: in vitro assembly behavior and filament architecture. J Biol Chem. 2013;288(6):4265–77. doi: 10.1074/jbc.M112.417030. PubMed PMID: 23204522; PMCID: PMC3567678.

62. Meijering E, Dzyubachyk O, Smal I. Methods for cell and particle tracking. Methods Enzymol. 2012;504:183–200. doi: 10.1016/B978-0-12-391857-4.00009-4. PubMed PMID: 22264535.

63. Upson RH, Haugland RP, Malekzadeh MN, Haugland RP. A spectrophotometric method to measure enzymatic activity in reactions that generate inorganic pyrophosphate. Anal Biochem. 1996;243(1):41–5. doi: 10.1006/abio.1996.0479. PubMed PMID: 8954523.

64. Carlier MF. Measurement of Pi dissociation from actin filaments following ATP hydrolysis using a linked enzyme assay. Biochem Biophys Res Commun. 1987;143(3):1069–75. doi: 10.1016/0006-291x(87)90361-5. PubMed PMID: 3566755.

65. Achanta SS, Varunan SM, Bhattacharyya S, Bhattacharyya MK. Characterization of Rad51 from apicomplexan parasite Toxoplasma gondii: an implication for inefficient gene targeting. PLoS One. 2012;7(7):e41925. doi: 10.1371/journal.pone.0041925. PubMed PMID: 22860032; PMCID: PMC3408395.

66. Firdaus ER, Park JH, Lee SK, Park Y, Cha GH, Han ET. 3D morphological and biophysical changes in a single tachyzoite and its infected cells using three-dimensional quantitative phase imaging. J Biophotonics. 2020;13(8):e202000055. doi: 10.1002/jbio.202000055. PubMed PMID: 32441392.

67. Suloway C, Pulokas J, Fellmann D, Cheng A, Guerra F, Quispe J, Stagg S, Potter CS, Carragher B. Automated molecular microscopy: the new Leginon system. J Struct Biol. 2005;151(1):41–60. doi: 10.1016/j.jsb.2005.03.010. PubMed PMID: 15890530.

68. Scheres SH. RELION: implementation of a Bayesian approach to cryo-EM structure determination. J Struct Biol. 2012;180(3):519–30. doi: 10.1016/j.jsb.2012.09.006. PubMed PMID: 23000701; PMCID: PMC3690530.

69. Zheng SQ, Palovcak E, Armache JP, Verba KA, Cheng Y, Agard DA. MotionCor2: anisotropic correction of beam-induced motion for improved cryo-electron microscopy. Nat Methods. 2017;14(4):331–2. doi: 10.1038/nmeth.4193. PubMed PMID: 28250466; PMCID: PMC5494038.

70. Rohou A, Grigorieff N. CTFFIND4: Fast and accurate defocus estimation from electron micrographs. J Struct Biol. 2015;192(2):216–21. doi: 10.1016/j.jsb.2015.08.008. PubMed PMID: 26278980; PMCID: PMC6760662.

71. Punjani A, Rubinstein JL, Fleet DJ, Brubaker MA. cryoSPARC: algorithms for rapid unsupervised cryo-EM structure determination. Nat Methods. 2017;14(3):290–6. doi: 10.1038/nmeth.4169. PubMed PMID: 28165473.

72. Dominguez R, Holmes KC. Actin structure and function. Annu Rev Biophys. 2011;40:169-86. doi: 10.1146/annurev-biophys-042910-155359. PubMed PMID: 21314430; PMCID: PMC3130349.

73. Terwilliger TC, Ludtke SJ, Read RJ, Adams PD, Afonine PV. Improvement of cryo-EM maps by density modification. Nat Methods. 2020;17(9):923–7. doi: 10.1038/s41592-020-0914-9. PubMed PMID: 32807957; PMCID: PMC7484085.

74. Croll TI. ISOLDE: a physically realistic environment for model building into low-resolution electron-density maps. Acta Crystallogr D Struct Biol. 2018;74(Pt 6):519–30. doi: 10.1107/S2059798318002425. PubMed PMID: 29872003; PMCID: PMC6096486.

75. Goddard TD, Huang CC, Meng EC, Pettersen EF, Couch GS, Morris JH, Ferrin TE. UCSF ChimeraX: Meeting modern challenges in visualization and analysis. Protein Sci. 2018;27(1):14–25. doi: 10.1002/pro.3235. PubMed PMID: 28710774; PMCID: PMC5734306.

76. Emsley P, Lohkamp B, Scott WG, Cowtan K. Features and development of Coot. Acta Crystallogr D Biol Crystallogr. 2010;66(Pt 4):486–501. doi: 10.1107/S0907444910007493. PubMed PMID: 20383002; PMCID: PMC2852313.

77. Afonine PV, Poon BK, Read RJ, Sobolev OV, Terwilliger TC, Urzhumtsev A, Adams PD. Real-space refinement in PHENIX for cryo-EM and crystallography. Acta Crystallogr D Struct Biol. 2018;74(Pt 6):531–44. doi: 10.1107/S2059798318006551. PubMed PMID: 29872004; PMCID: PMC6096492.

78. Adams PD, Afonine PV, Bunkoczi G, Chen VB, Davis IW, Echols N, Headd JJ, Hung LW, Kapral GJ, Grosse-Kunstleve RW, McCoy AJ, Moriarty NW, Oeffner R, Read RJ, Richardson DC, Richardson JS, Terwilliger TC, Zwart PH. PHENIX: a comprehensive Python-based system for macromolecular structure solution. Acta Crystallogr D Biol Crystallogr. 2010;66(Pt 2):213–21. doi: 10.1107/S0907444909052925. PubMed PMID: 20124702; PMCID: PMC2815670.

79. Pollard TD, Cooper JA. Actin and actin-binding proteins. A critical evaluation of mechanisms and functions. Annu Rev Biochem. 1986;55:987–1035. doi: 10.1146/annurev.bi.55.070186.005011. PubMed PMID: 3527055.

